# Rapid genome editing by CRISPR-Cas9-POLD3 fusion

**DOI:** 10.1101/2021.05.23.445089

**Authors:** Ganna Reint, Zhuokun Li, Kornel Labun, Salla Keskitalo, Inkeri Soppa, Katariina Mamia, Eero Tölö, Monika Szymanska, Leonardo A. Meza-Zepeda, Susanne Lorenz, Artur Cieslar-Pobuda, Xian Hu, Diana L. Bordin, Judith Staerk, Eivind Valen, Bernhard Schmierer, Markku Varjosalo, Jussi Taipale, Emma Haapaniemi

## Abstract

Precision CRISPR gene editing relies on the cellular homology-directed DNA repair (HDR) to introduce custom DNA sequences to target sites. The HDR editing efficiency varies between cell types and genomic sites, and the sources of this variation are incompletely understood. Here, we have studied the effect of 450 DNA repair protein - Cas9 fusions on CRISPR genome editing outcomes. We find the majority of fusions to improve precision genome editing only modestly in a locus- and cell-type specific manner. We identify Cas9-POLD3 fusion that enhances editing by speeding up the initiation of DNA repair. We conclude that while DNA repair protein fusions to Cas9 can improve HDR CRISPR editing, most need to be optimized to the particular cell type and genomic site, highlighting the diversity of factors contributing to locus-specific genome editing outcomes.

## Introduction

CRISPR/Cas9 has become a common genome editing tool in both basic and medical research. It induces targeted DNA double-strand breaks (DSBs) that are commonly repaired by non-homologous end-joining (NHEJ), which leads to gene disruption and knockout. The less common pathways utilize homology-directed repair (HDR), which can induce custom genetic changes to the DSB sites by using exogenous DNA as a template. HDR is confined to the synthesis (S) phase of the cell cycle^1^. Therefore, its efficiency varies between cell types^2^ and improves upon rapid cell proliferation^3^ and upon targeting the Cas9 expression to S cell cycle phase^4^. Factors that impair cell cycle progression from G1 to S decrease CRISPR-Cas9 mediated HDR^5^. HDR can be promoted by increasing the local concentration of the repair template^6^ and by general NHEJ pathway inhibition^7^,^8^,^9^, which, however, often negatively affects the cell viability and fitness^10^,^11^. Overall, genome editing is most efficient in open chromatin^12^ and when Cas9 is bound to an actively transcribed DNA strand where the approaching polymerase can rapidly remove the complex^13^,^14^,^15^. Cas9 binds to the DNA for extended periods^14^,^15^ and successful cut/repair requires the activation of pathways typically involved in resolving stalled DNA replication forks^16^.

Enhancing CRISPR-based HDR by Cas9 fusion proteins can stimulate correction locally at the cut site, without causing generalized disturbance in the cellular DNA repair process and thus increasing the safety and specificity of the editing. HDR-based precision genome editing can be enhanced by fusing Cas9 with DNA repair proteins or their parts^17^,^18^, chromatin modulating peptides^19^ or peptides that locally decrease NHEJ pathway activation at the DNA double strand break sites^20^. The effect of the published fusions has been guide- and locus-specific, in line with the fact that CRISPR-Cas9 editing efficiency depends heavily on the target sequence.

Here we systematically characterize the effect of ~450 human DNA repair protein and protein fragment fusions on CRISPR-Cas9-based HDR genome editing, using reporter human embryonic kidney (HEK293T) and immortalized retinal pigment epithelium (RPE1) cell models. We note that for most of the proteins, their effect on the HDR editing of the reporter locus is modest. For the subset that improve genome editing, the effect is locus-specific and comparable to previous publications that have studied single Cas9 fusions ^18^. We identify Cas9 fusion to DNA polymerase delta subunit 3 (Cas9-POLD3) as a new HDR enhancer. The fusion enhances editing at the early time points by speeding up the kinetics of Cas9 DNA binding and dissociation, allowing the rapid initiation of DNA repair.

## Results

### The effect of DNA repair protein fusions on CRISPR-Cas9 genome editing

We sought to better understand the processes that govern the choice and efficiency of DNA repair in CRISPR-Cas9 induced breaks. To this end, we conducted arrayed screens where we fused ~450 human proteins and protein domains involved in DNA repair to the C-terminus of wildtype *S. pyogenes* Cas9 (Cas9WT, Fig. 1A-B, Fig. S1 Supplementary file DNA repair domain library.xlsx). The reporter cell line contains a non-functional Green Fluorescent Protein (GFP), a guide targeting the mutant GFP (sgGFP), and functional Blue Fluorescent Protein (BFP) as a positive selection marker. The proportion of cells that turned GFP positive was used as a proxy for HDR efficiency. After the initial screen, we chose to validate Cas9WT fusions with >4% GFP+ cells per well or HDR improvement >2 times relative to experiment average (total 52 fusions from two independent screens) (Fig. 1B and Online Methods). We transfected the fusions to clonal reporter HEK293T and RPE1 (hTERT immortalized retinal pigment epithelium) cell lines. RPE1 has a near-normal karyotype and is the target cell type in eye gene therapy.

**Figure 1.**
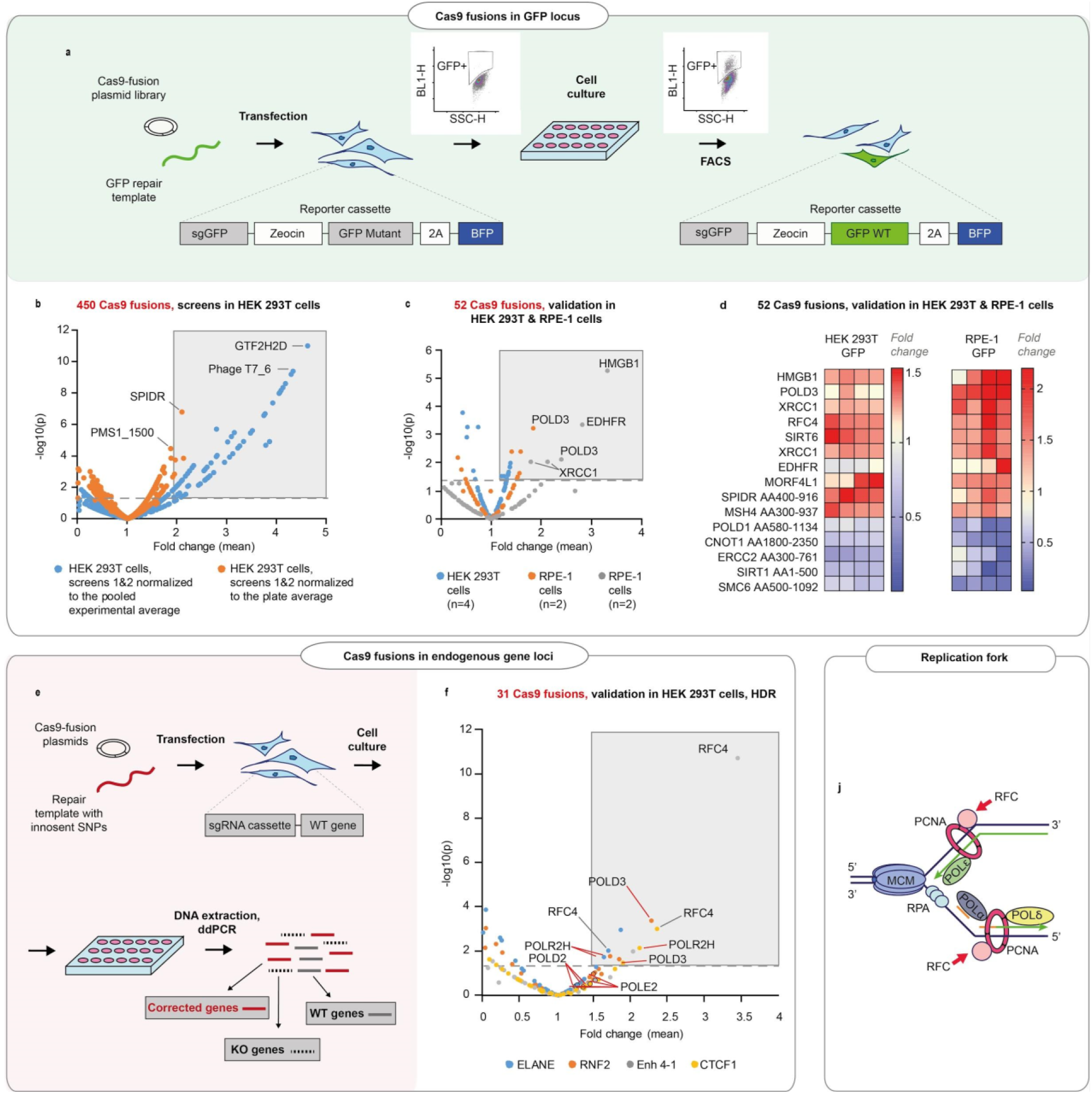
DNA repair proteins that affect genome editing outcomes. **a**. Schematic representation of the screen. HEK293T cell line contains a reporter cassette expressing a selection marker, guide (sgGFP) and a mutant GFP-BFP protein. The line is transfected with a GFP repair oligo and an arrayed plasmid library containing ~450 DNA repair proteins that are fused to the C-terminus of Cas9WT. The HDR editing efficiency for each fusion is defined by the percentage of cells with restored GFP function in each transfected well (measured by FACS). **b.** Normalized GFP recovery values from two independent screens, both experiments n=3, each data point represents an average of value of all six replicas (n=2×3), where each replicate was normalized either to the experimental average (blue dots) or to the plate average (orange dots). P-values were calculated by one-way Anova test, where the mean of each triplicate is compared to the combined mean of all other triplicates from the screen. **c.** Experiment average-normalized GFP recovery values for 52 fusions, chosen based on their performance in panel (b). In HEK293T n=4, one independent biological experiment, in RPE1 n=2, two independent biological experiments. Statistical significance is calculated as in (b). **d.** Normalized GFP recovery values for ten best-performing and five worst-performing Cas9 fusions from panel (c). Protein fragments are denoted based on the position in canonical transcript (ie. SPIDR AA400-916 corresponds to a fragment of SPIDR protein which starts at amino acid 400 and ends at amino acid 916). **e.** Schematic representation of the experiment shown in f. **f.** Experiment average - normalized HDR editing for 31 fusions in four endogenous loci (ELANE, RNF2, Enh 4-1 and CTCF1) in HEK293T cells, quantified by droplet digital PCR. Polymerase fusions are marked with red pointers. n=3, one independent biological experiment for each locus. Statistical significance is calculated as in (b). The raw data points are visible in Fig. S3. **j.** Replication fork schematics.

We found that in the GFP reporter locus, the proteins that improved HDR above the experiment mean were largely similar between RPE1 and HEK293T cells (Fig 1C-D and Fig S2A). For 31 fusions that demonstrated a stable performance in validation experiments, we also quantified the HDR and NHEJ editing by droplet digital PCR (ddPCR) in three transcribed and one non-transcribed endogenous loci in HEK293T cells (Fig 1E-F and Fig S2B, Fig S3). We discovered a number of diverse polymerases among the better-performing fusions (POLD3, POLD2, POLR2H, POLE2) (Fig S2B, Fig S3). In addition, the replication factor C subunits RFC4 and RFC5 performed well in most of the tested loci. POLD3, POLE2, RFC4 and RFC5 are all members of the DNA replication fork machinery (Fig. 2J).

**Figure 2.**
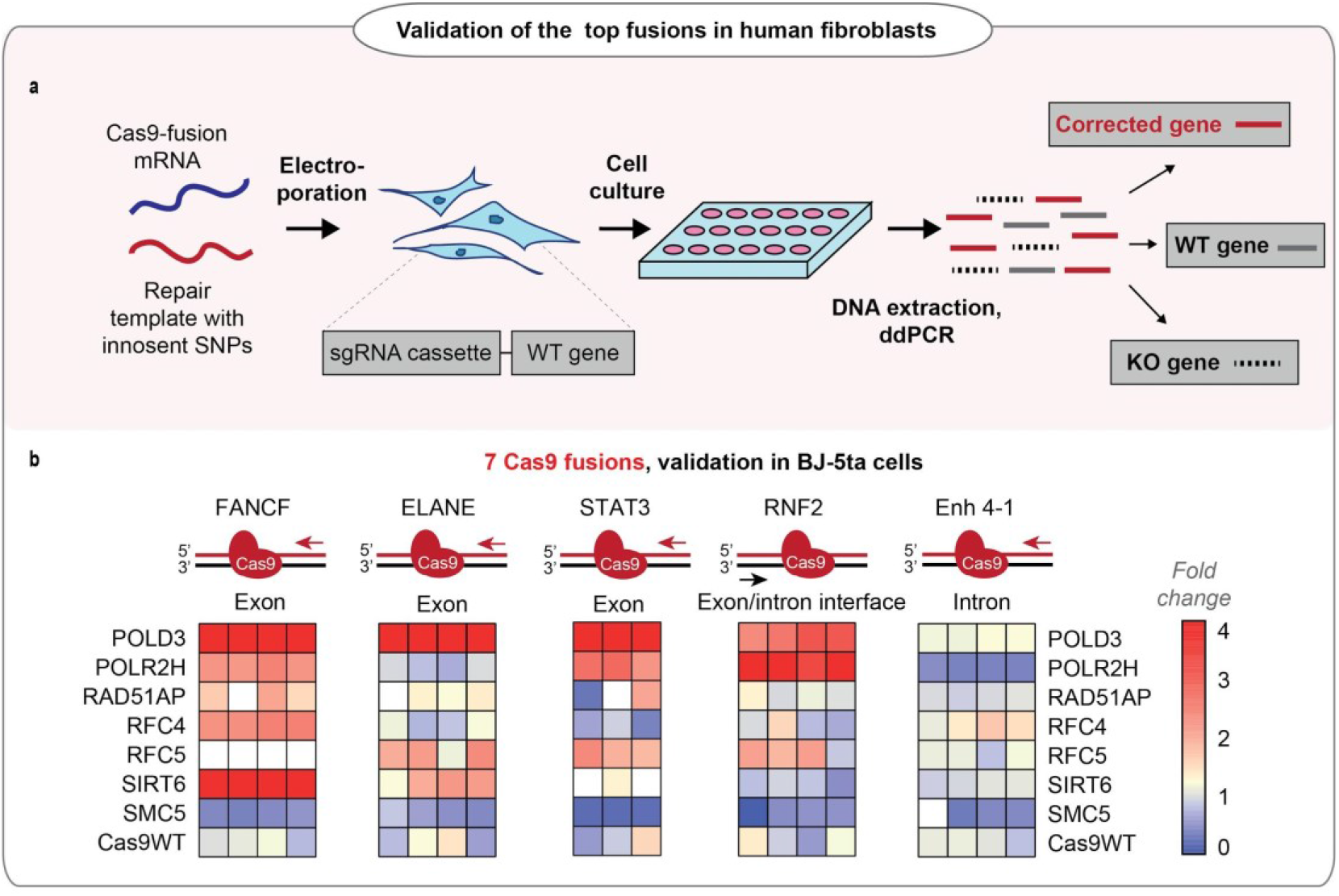
Performance of the fusions in BJ-5ta cell line. **a.** Schematic representation of the experiment shown in b. **b.** Normalized HDR efficiency of the seven bestperforming fusions in five endogenous loci in hTERT immortalized fibroblasts (Bj5-ta), quantified by ddPCR. Heat maps represent values normalized to Cas9WT for each gene locus. The sets have an upper limit threshold and values above it are color-coded as a scale maximum (bright-red). One independent experiment for each gene locus, n=4 for FANCF, Enh 4-1, RNF2, ELANE gene loci; n=3 for STAT3 gene. The raw data points are visible in Fig. S4.

We chose seven Cas9 fusions for the final validation, and transfected them as mRNA in human hTERT immortalized fibroblasts (BJ-5ta) using five endogenous loci as models (Fig. 2A-B, Fig. S4). The fusion performance was strongly dependent on individual loci, with POLD3 outperforming the other fusions across the majority of tested conditions (Fig. 2B, Fig. S4). We further tested the performance of Cas9-POLD3 across decreasing concentrations in the RPE1 reporter cells and noted a twice-fold, concentration-independent benefit to Cas9WT (Fig. S5).

### Cas9-POLD3 fusion accelerates the initiation of cut repair

POLD3 is a component of the replicative polymerase **δ**, and we hypothesized that POLD3 fusion improves genome editing by speeding up the removal of Cas9 from the cut site^13^. The faster Cas9 removal results in early recruitment of the DNA damage response machinery to the break and speeds up the DNA repair progression.

To understand whether the speed of the overall DSB DNA repair is increased after Cas9-POLD3 editing, we quantified the HDR and NHEJ repair across time for eight fusions in the RPE1 GFP reporter locus by ddPCR (Fig. 3A,B). For Cas9WT and the other fusions, there was a linear increase in successful editing towards later (72h) timepoints. For Cas9-POLD3, the editing peaked earlier (24h) and plateaued at the later (72h) timepoint. To further quantify how rapidly the DNA damage response activates upon cutting the DNA, we compared the emergence of the early-stage DNA damage marker γH2AX^21^ which arrives at DNA double-strand breaks within minutes after CRISPR cutting (Fig. 3C-E). We transfected BJ5-ta fibroblasts with Cas9WT or Cas9-POLD3 mRNA and a pool of 4 guides targeting three coding (RNF2, STAT3, Enh4-1) and one non-coding locus (CTCF1) and quantified the γH2AX foci with immunofluorescence microscopy. For Cas9WT, the number of foci steadily increased and peaked at 24h, after which the count declined. For Cas9-POLD3 the number of foci was already high in the early timepoints (Fig. 3E), suggesting rapid Cas9-POLD3 recruitment and removal from the DNA. In parallel to γH2AX foci counting, we quantified the NHEJ progression across time by ddPCR in each individual locus (Fig. 3F-I). We saw a profoundly increased NHEJ in Cas9-POLD3 treated samples in early (8-12h) timepoints, with the exact change depending on the locus and becoming less distinct after 24 hours. All these experiments suggest that the mechanism for improved gene editing for Cas9-POLD3 is the rapid initiation of DNA repair, possibly due to the early recruitment and removal of the fusion from the DNA. The effect size is locus dependent and is not evident in other fusions where the editing follows the kinetics of Cas9WT.

**Figure 3.**
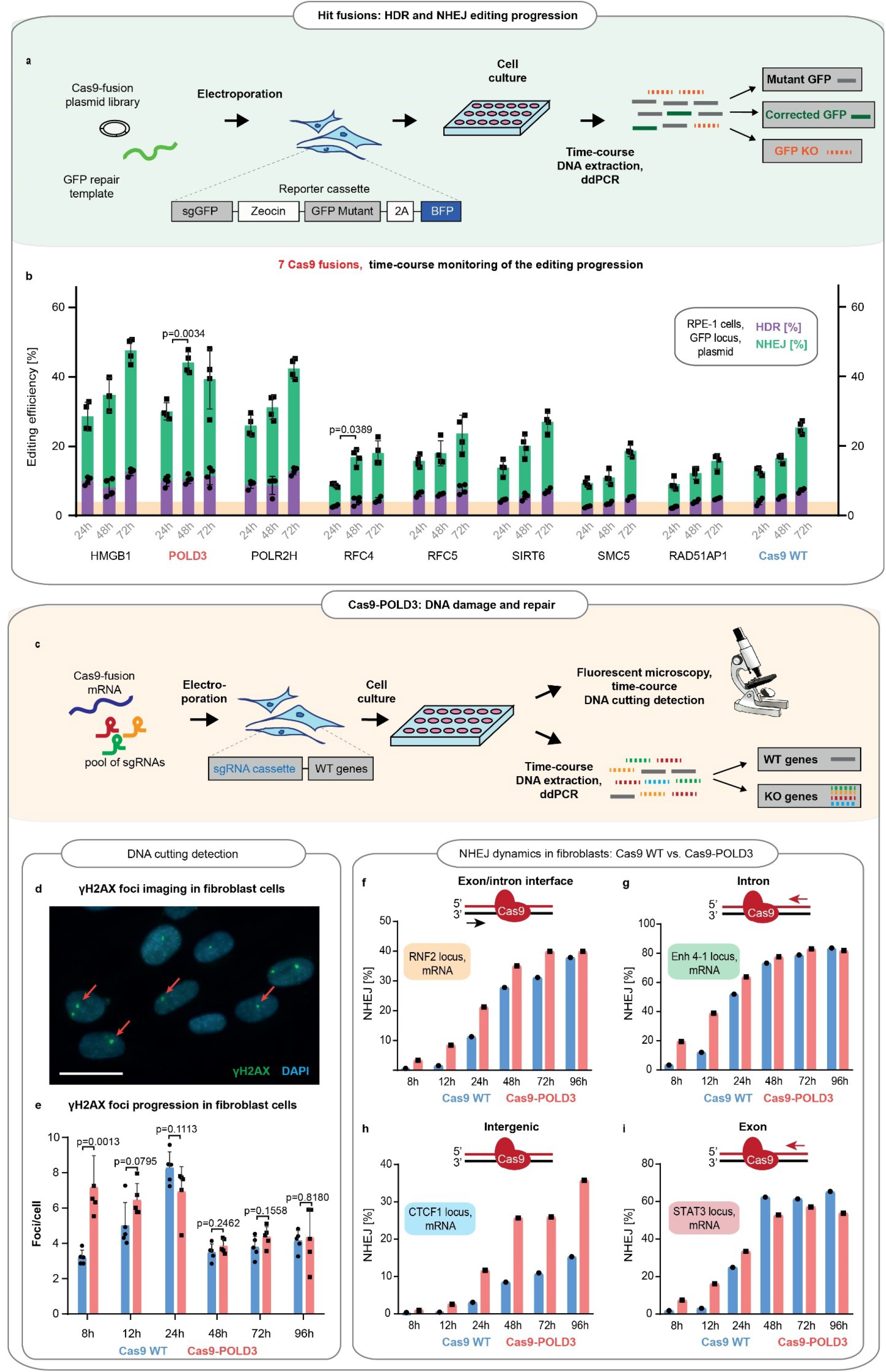
Comparison of the DNA repair dynamics between Cas9 fusions. **a.** Schematic representation of experiment shown in b**. b**. The DNA repair progression in the RPE-1 reporter locus for eight of the better-performing fusions (selected from screens on Fig. 1D-E). ddPCR quantification, n=3, single independent experiment, bar denotes mean value, error bars represent ± S.D. Orange highlighting indicates the Cas9WT HDR editing level at 24h post-electroporation. P-values denote significance of the total editing (HDR+NHEJ) increment between the Cas9WT and other fusions (for 24h to 48h period). Statistical values derived using one-way ANOVA test. **c.** Schematic representation of experiment shown in d-i. **d.** Representative immunofluorescence image used for DNA breakpoint quantification. Cell nuclei are depicted in blue, γH2AX foci in green, and red arrows indicate individual foci. Scale bar 50 um. **e.** Time course for γH2AX foci emergence in human immortalized fibroblasts (BJ5-ta). The cells were electroporated with Cas9WT or Cas9-POLD3 mRNA and a pool of three guides targeting RNF2, Enh4-1, and STAT3 loci. The CTCF1 gRNA is constitutively expressed. Five confocal images were taken for each condition, n=5, single independent experiment, bar denotes mean value, error bars represent ± S.D. Statistical significance of the difference between Cas9WT and Cas9-POLD3 is calculated using ANOVA test for the equality of the means at a particular time point. **f-i.** ddPCR quantification of the NHEJ repair dynamics across time for the experiment described in e. The red arrow shows the direction of the approaching polymerase. The DNA strand which the CRISPR-Cas9 binds to is colored in red. n=1, single independent experiment, bar denotes mean value, error bars represent ± S.D.

### Cas9-POLD3 interacts with ATP-dependent helicases

To better understand how the fusion proteins modify editing, we looked at their formed molecular interactions (interactomes) using affinity-purification mass spectrometry (AP-MS) (Fig. 4A-B, Table S1). We first constructed HEK293T cell lines consisting of the GFP reporter and tet-inducible, MAC-tagged^22^ Cas9 fusions in the isogenic flip-in^®^ locus (Fig. S6). When Cas9 expression is induced, it complexes with the GFP targeting guide and binds and cuts the GFP sequence. The MAC-tag in the Cas9 ribonucleoprotein (RNP) complex enables the affinity purification of the complex. Additionally, the MAC-tag enables proximity-dependent biotinylation of the interacting and close-proximity proteins that can be subsequently purified and identified by mass spectrometry (BioID labelling).

**Figure 4.**
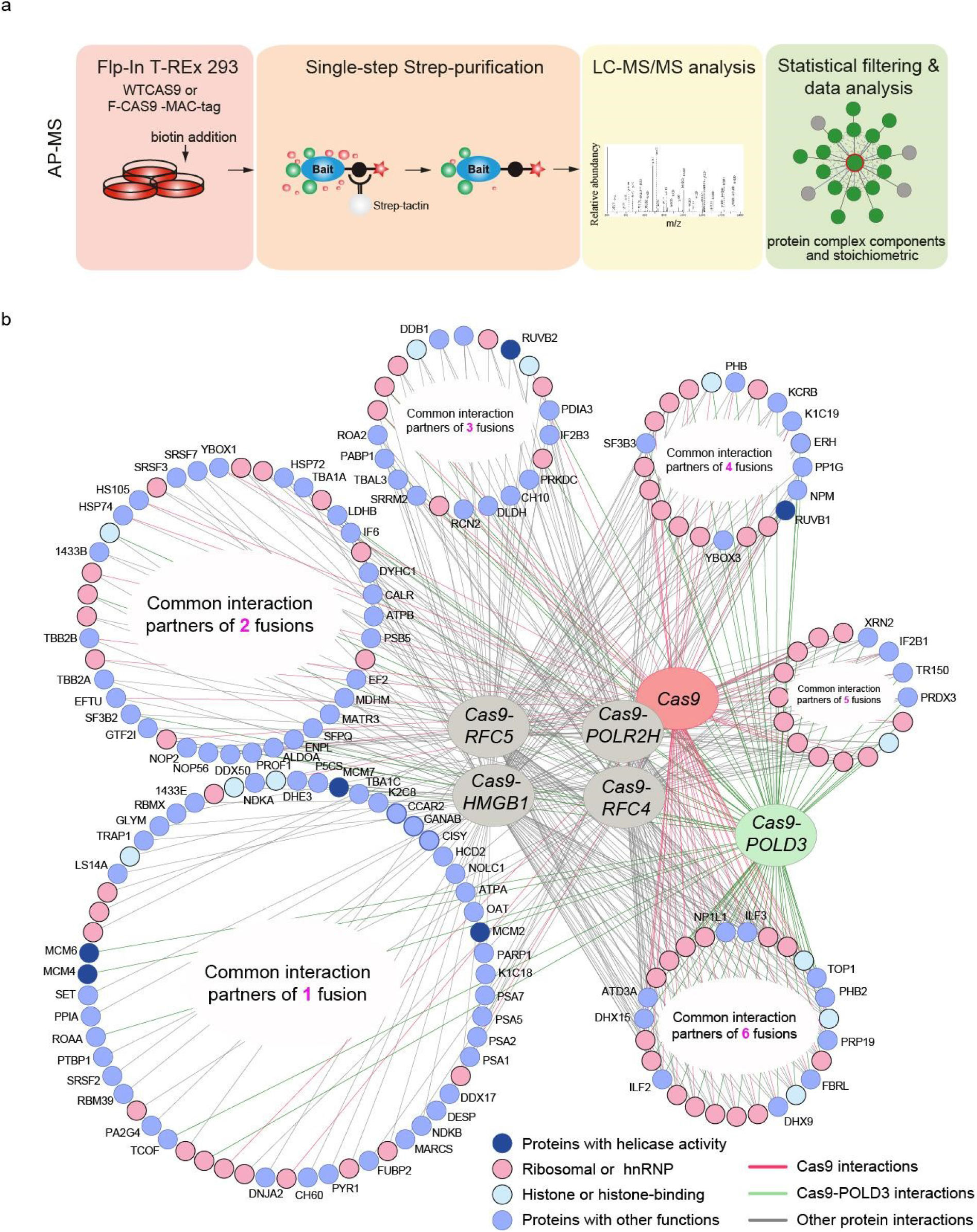
Protein-protein interactions of editing-enhancing Cas9 fusions. **a.** Schematic representation of the Affinity Purification Mass Spectrometry (AP-MS) experimental workflow. **b.** Protein interaction map of the Cas9 fusions (three biological replicates). The interacting proteins are clustered based on the number of interactions they make with the Cas9 fusions (I, unique interaction partners, VI, interact with all Cas9 fusions, including Cas9WT). Pink circles represent RNA-binding proteins, light-blue circles represent histone proteins, dark blue circles represent interaction partners with helicase activity, and light blue circles denote proteins with other functions.

We mapped the stable interactions for Cas9WT and five Cas9 fusions. These include three DNA replication fork proteins (POLD3, RFC4 and RFC5), the RNA polymerase POLR2H and HMGB1, which is a structural chromatin component and a nucleosome remodeler. We recovered a total of 76 Cas9-POLD3 interactions, none of which have been previously reported (there are a total of 9 reported POLD3 interactions). We did not recover any polymerase δ subunit components, likely because the fusion uses the short POLD3 isoform 3 (ENST00000532497.5) which lacks the amino acids 1-106 that interact with other polymerase δ subunits ^23^. Cas9-RFC4 and Cas9-RFC5 had one known interactor for each (both RFC4 and RFC5 have >40 reported interactions). Cas9-HMGB1 had 103 interactors (29 previously reported for HMGB1). The landscape for the Cas9 fusions thus differs somewhat from their corresponding endogenous proteins, particularly for the replication fork components.

Cas9WT and all of the fusions recovered a large number of RNA-binding proteins that presumably bind to the guide, as well as histones and other structural DNA components. The unique interactions of the Cas9 fusions included proteins with known nucleosome remodeling and helicase activity (Table S2). These include the minichromosome maintenance protein complex (MCM) that interacts with POLD3. Other helicases were RUVB1 and 2 that interacted with the other fusions. MCM and RUVB1-2 are helicase complexes that participate in DNA repair and replication fork stability maintenance^24^,^25^. The helicases might aid editing by opening DNA and making it more accessible to Cas9 binding. After cutting, they might dislodge Cas9 from the cutting site and make the DNA ends available for processing by the DNA repair machinery^14^,^15^,^13^.

### Cas9-POLD3 fusion does not increase off-target cuts

The POLD3 fusion alters the DNA binding kinetics of the CRISPR complex and might affect the specificity of the CRISPR binding and cutting. Therefore, we compared the off-target profiles of the Cas9WT and Cas9-POLD3 using GUIDE-Seq^26^. Briefly, we electroporated HEK293T cells with plasmids encoding the fusion, the sgRNA and blunt phosphorylated double-stranded oligodeoxynucleotides (dsODN), which incorporate in the DNA double-strand breaks and are used for selective amplification and sequencing of the DNA break sites. We used a previously published guide ^26^ against HEK site 4, which is a non-coding target in the HEK293T genome. The cutting with this guide leads to a large number of off-target events, with several off-target sequences being cut more frequently than the on-target site ^26^.

The off-target signature of the Cas9-POLD3 was largely overlapping with the wild type nuclease, with slightly less cut sites for Cas9-POLD3 (Fig. 5A-B), suggesting that the polymerase fusion does not alter the cutting specificity of Cas9. We also compared the indel profiles of Cas9WT and Cas9-POLD3 by deep sequencing of the RNF2 and ELANE target loci (Fig. 5C-F). We used Unique Molecular Identifiers (UMIs) in the amplicon primers to account for potential PCR bias. The indel profiles were dependent on the target sequence, with NHEJ (1bp deletion) dominating in RNF2 and MMEJ (23bp deletion) dominating in ELANE locus. The outcomes were generally similar between Cas9WT and Cas9-POLD3. Based on these experiments, the polymerase fusion does not affect the CRISPR on- or off-target repair outcomes. However, a larger experiment is necessary for a more comprehensive map of the specificity and the repair outcomes for Cas9-POLD3.

**Figure 5.**
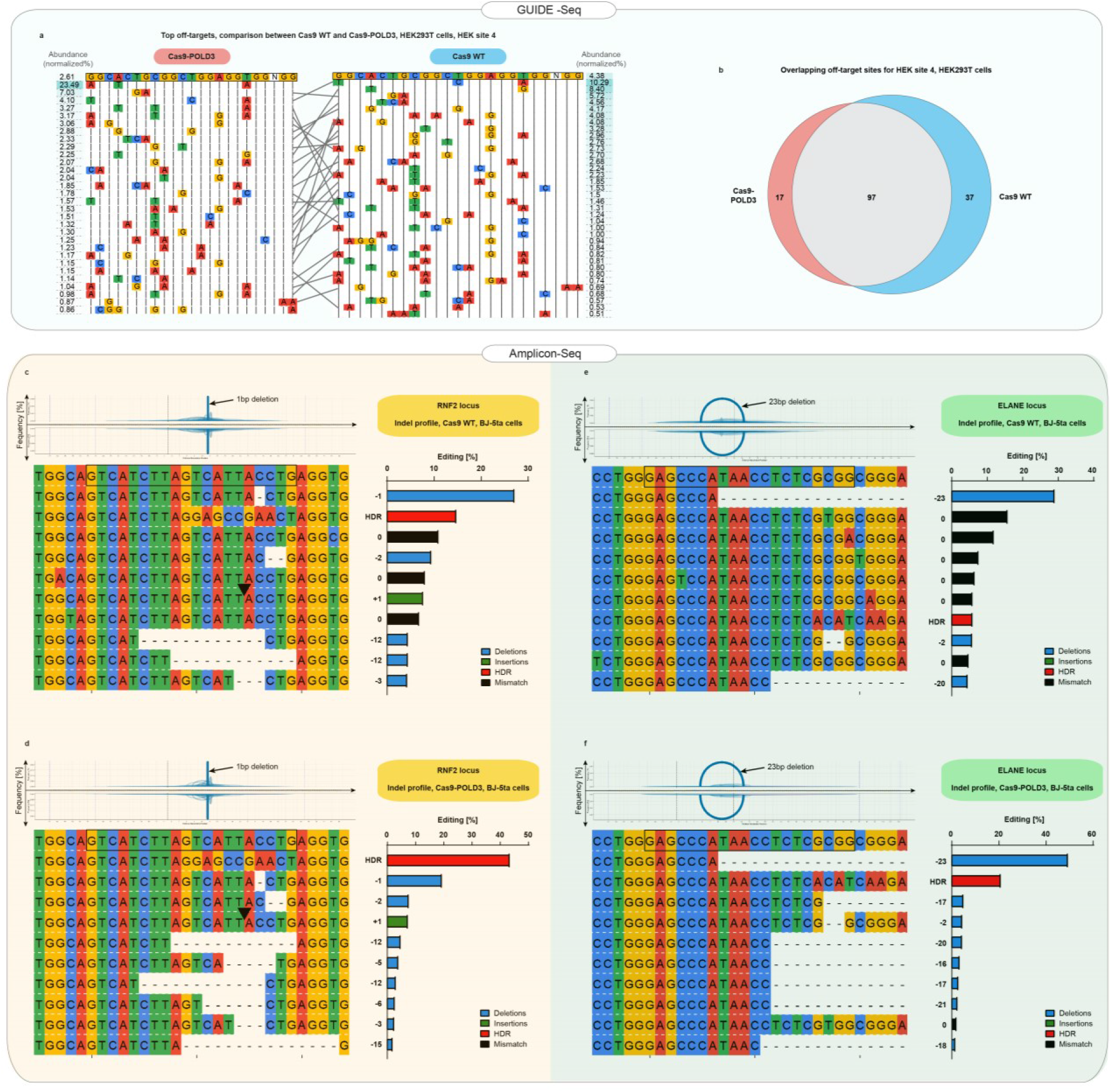
Off-target and indel profiles of WT and Cas9-POLD3 nucleases. **a.** Mismatch plot of the Guide-seq^6^ results for Cas9WT and Cas9-POLD3, using a published guide^6^ against the endogenous HEK site 4. The on-target sequence is depicted at the first line of the table. The most abundant off-targets are listed underneath the intended edit and sorted by frequency, which is determined by dividing the number of reads that contain the off-target edit with the total read count. Gray lines connect the shared off-target sites. **b.** Venn diagram of the common and unique off-target sites between Cas9WT and Cas9-POLD3. **c-f.** Mismatch plots of the indel profiles of Cas9WT and Cas9-POLD3 in BJ-5ta fibroblasts, obtained by deep amplicon sequencing. The on-target gRNA binding site is on the top row, highlighted with a black box. The most frequent indels are below, sorted according to their frequency. The frequency is calculated by dividing the indel read count by the sum of the top 10 read counts. HDR template-matching edits marked with the red on the summary plots. Plots depict: **c.** On-target editing signature of Cas9 WT, RNF2 locus. **d.** On-target editing signature of Cas9-POLD3, RNF2 locus. **e.** On-target editing signature of Cas9 WT, ELANE locus. **f.** On-target editing signature of Cas9-POLD3, ELANE locus.

### Cas9-POLD3 fusion comparison to published improvements

Several Cas9 fusions have previously been reported to increase HDR editing^18^,^20^,^4^. Of the published fusions, our screening independently identified Cas9-HMGB1^19^ as an editing enhancer. For a broader comparison, we evaluated the performance of Cas9-POLD3 against five additional published fusions: Cas9-DN1S (fragment inhibiting NHEJ initiation by 53BP1 blockade)^20^, Cas9-CtIP (a protein that promotes HDR pathway choice)^18^, Cas9-HE (active fragment of CtIP)^18^, Cas9-Geminin (fragment that targets the Cas9 expression to S-G2 phases of the cell cycle)^4^ and Cas9 HE-Geminin (the fusion combines both HE and Geminin fragments)^18^. The five fusions showed comparable performance with Cas9-POLD3, with S cell cycle phase targeting providing the highest benefit (Fig. 6A). To understand whether it was possible to use the fusions as CRISPR-Cas9 ribonucleoprotein complexes, we produced Cas9 fusions to POLD3, HMGB1 and HMGB2 in *E.coli* and transfected them with cationic lipid transfer to RPE-1 and HEK293T reporter cell lines. We excluded Cas9-Geminin, as it has previously been tested as an RNP complex^27^. Cas9-POLD3 performed best in both cell lines, although we cannot exclude differences in transfection efficiency and fusion protein folding to affect editing outcomes (Fig. 6B). We also tested the effect of Cas9-POLD3 fusion activity in human embryonic stem cells line and stimulated peripheral blood mononuclear cells (Fig. S7). Although the POLD3 fusion showed a trend for improvement in both cell types, the benefit is less pronounced than in the model RPE1 and fibroblast cells, possibly due to cell-type specific differences in DNA damage and repair signaling.

**Figure 6.**
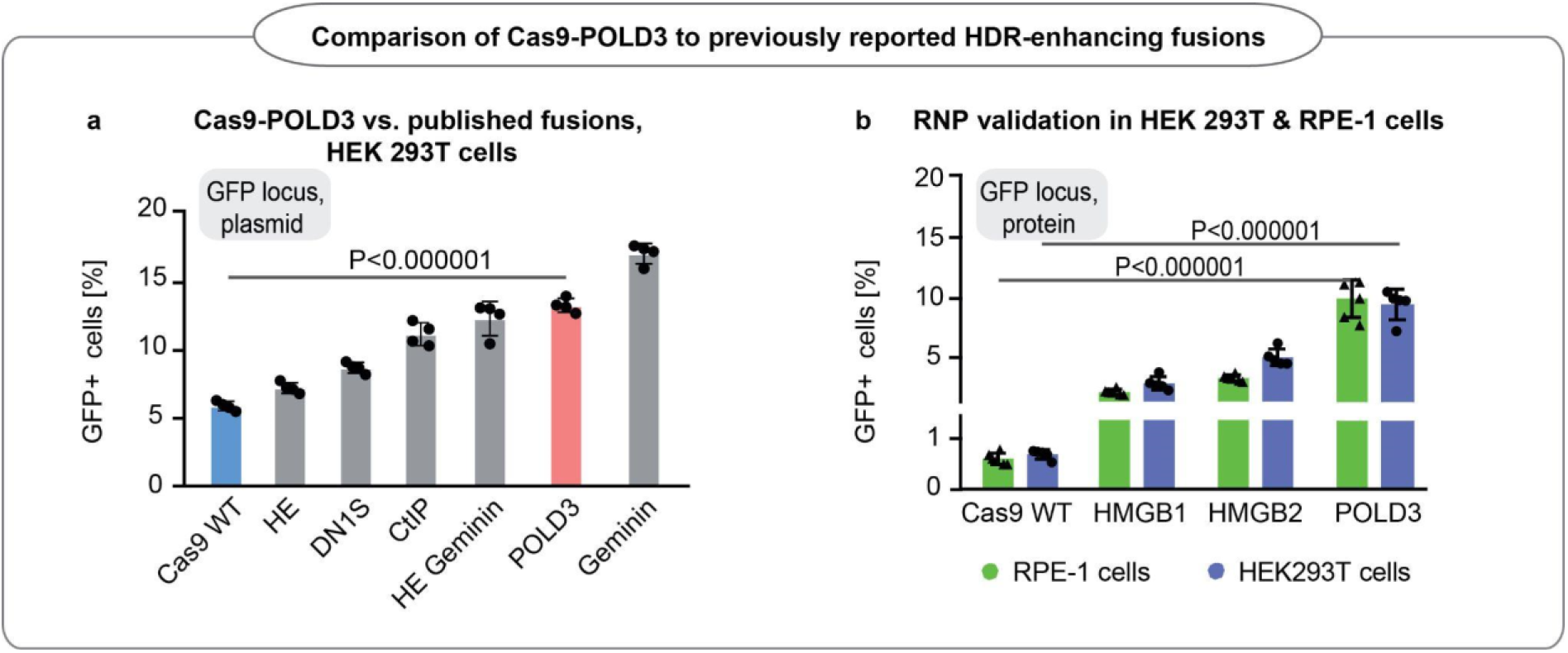
Editing efficiency of Cas9-POLD3 and a panel of various HDR improving fusions reported in the literature. **a**. Editing of GFP in reporter HEK293T cells. n=4, one of two independent experiments, bar denotes mean value, error bars represent ± S.D. Statistical significance is calculated with unpaired, two-sided Student’s t-test. **b.** GFP reporter locus editing efficiency of recombinant Cas9 fusion proteins in RPE-1 and HEK293T cells. n=5, one of two independent experiments, bar denotes mean value, error bars represent ± S.D. Statistical significance is calculated with Anova that is adjusted for multiple comparisons.

## Discussion

Through screening 450 human proteins involved in DNA repair, we have identified a subset that can improve genome editing as WT Cas9 fusions. The editing-enhancing fusions show locus- and cell type specific effects^18^. We have further identified a Cas9 fusion to POLD3, which improves editing by hastening the kinetics of DNA repair.

DNA polymerase delta subunit 3 (POLD3) is a component of the replicative polymerase δ, an enzyme that is involved in DNA replication and repair^28^, ^29^, ^30^. We show that the POLD3 fusion to Cas9 accelerates the initiation of the CRISPR cut repair. The CRISPR complex binds to DNA for 6-8 hours after cutting and needs active removal from the DNA prior to the start of the repair. Approaching RNA polymerase can dislodge Cas9 from the DNA break, which exposes the cut strands to the DNA repair machinery and speeds up the repair process^13^. We hypothesize that POLD3 fusion partially shares this mechanism and improves genome editing by accelerating the removal of Cas9 from the cut site^13^. The faster Cas9 removal results in early recruitment of the DNA damage response machinery to the break, which speeds up the DNA repair progression. The final HDR improvement depends on the individual locus and is independent from the status of the RNA transcription on the edited strand. It is unlikely that Cas9-POLD3 locally promotes DNA replication, as the indel and off-target profile is unaltered between Cas9WT and Cas9-POLD3.

Cas9 fusions can stimulate homologous recombination in a number of ways, including timing the editing to S cell cycle phase^4^ improving chromatin accessibility^19^, or altering the local balance of DNA repair factors ^20^, ^18^. The published strategies provided approximately two-fold editing improvement, when tested as plasmids in our reporter HEK293T line. This level is comparable to other editing-enhancing fusions that we identified in our screen. In addition to altering the DNA repair pathway choice, it is possible to improve editing by increasing the target sequence accessibility through chromatin remodeling ^19^. Our screen identifies two nucleosome remodeling enzymes (SIRT6 & HMGB1) among the better-performing Cas9 fusions; of these, HMGB1 has previously been reported as editing-enhancing^19^. In comparison to the previously reported fusions, POLD3 provides a novel mechanism for editing enhancement, and combining the different strategies would likely increase the overall efficacy.

The efficiency of CRISPR cutting and the subsequent DNA repair is dependent on the individual locus, where the cell type, guide sequence, chromatin state, the DNA strand orientation and the level of the locus transcription all affect the editing outcome ^31^, ^32^. Similarly, in this study we note locus- and cell-type specific editing enhancement for the screened fusions. There is a complex and poorly understood interplay between the CRISPR-Cas9 complex and the local genomic landscape, and it can be challenging to find CRISPR-Cas9 fusions that promote HDR across variable conditions.

In conclusion, we have extensively compared the known human DNA repair proteins and protein domains for their ability to locally improve CRISPR-Cas9 genome editing. We confirm that rapid removal of the CRISPR complex from the cut site and improved chromatin accessibility can locally promote HDR outcomes. We also note that the effects are locus- and cell type specific, suggesting that the strategy needs to be tailored to individual applications.

## Supporting information

Supplementary Table 1_AP-MS_BioID_Cas9

Supplementary Table 5_Statistics

DNA_repair_domain_library

## Acknowledgements

Norwegian Research Council, the South-Eastern Norway Regional Health Authority, Knut and Alice Wallenberg Foundation, Cancerfonden, Barncancerfonden, Instrumentarium Foundation and Academy of Finland supported this work. Part of this work was carried out at the High Throughput Genome Engineering Facility and the Swedish National Genomics Infrastructure funded by Science for Life Laboratory. We would like to thank the Protein Science Facility at Karolinska Institutet for recombinant Cas9 production, and the HSØ Genomics Core Facility at Oslo University Hospital for providing the high-throughput sequencing. Sini Nieminen is acknowledged for expert technical assistance. We thank Alicia Roig-Merino and Caoimhe Nic An tSaoir for their advice on MaxCyte electroporation experimental design.

## Author contributions

G.R. and E.H. wrote the manuscript. E.H. and B.S. conducted the initial DNA repair protein screens. G.R., Z.L., I.S. and K.M. performed the subsequent validation experiments. A.C.P. and J.S. provided hESC cells and maintained the cells throughout experiments. X.H and D.L.B. contributed to establishment of the γH2AX foci counting pipeline. L.M. and S.L provided help in High Throughput Sequencing. K.L and E. V. performed the sequencing data analysis. E.T. provided expertise in data analysis. S.K. and M.V. performed the protein interactome profiling. E.H. and J.T. supervised the study. All authors read and approved the final manuscript.

## ONLINE METHODS

### Cell culture

HEK293T, RPE1 and BJ-5ta cells were cultured at 37°C in a humidified incubator in DMEM (Thermo Fisher Scientific) supplemented with 10% fetal bovine serum (FBS; Thermo Fisher Scientific) and 1 % Penicillin-Streptomycin (Thermo Fisher Scientific). After electroporation, cells were cultured in DMEM (Thermo Fisher Scientific) supplemented with 10% FBS (Thermo Fisher Scientific), but without penicillin-streptomycin. Cells were split routinely twice a week using TrypLE^™^ Express Enzyme (Thermo Fisher Scientific).

### RPE-1 and HEK293T GFP reporter cell lines

For easy detection of CRISPR-Cas9-mediated homologous recombination, we created a cassette encoding a) Zeocin resistance gene for selecting stable integrants, b) mutant Green Fluorescent Protein (GFP) (37A-38A-39A) sequence, c) Blue Fluorescent protein (BFP) or Red Fluorescent protein (RFP) sequence separated from GFP by a 2A self-cleaving peptide and c) sgRNA targeting the mutant GFP. When Cas9 along with a repair template correcting the GFP mutation is introduced into these cell lines, gene correction will give rise to functional GFP. GFP fluorescence was measured by FACS after 5 days if Cas9 was delivered as a plasmid, or 4 days if Cas9 was delivered as ribonucleoprotein.

The reporter cassette sequence was cloned to pLenti sgRNA(MS2) plasmid backbone (Addgene #61427). The cassette sequence is shown in Supplementary Appendix III and the plasmid will be deposited to Addgene.

### Lentivirus production and transduction

The reporter cassette was packaged into lentiviral particles by transfecting HEK 293T cells with the reporter cassette plasmid and the two packaging plasmids psPAX2 (Addgene #12260) and pCMV-VSV-G (Addgene # 8454) in equimolar ratios. After 48 hours, the virus-containing supernatant was concentrated 40-fold using Lenti-X concentrator (Clontech). Single use aliquots were prepared and stored at −140°C.

The cassette was transduced to HEK293T and RPE-1 cells by lentivirus at low multiplicity of infection (MOI). The cells were propagated in Zeocin (500 μg/ml) selection media for two weeks. After selection, single-cell clones were obtained by sorting BFP+ cells to 96-well plates by flow cytometry. The clones were expanded and frozen for future use.

### BJ-5ta and HEK293T sgRNA cell lines

The cell lines express a guide targeting one of the six endogenous loci: *RNF2*, *ELANE*, *Enh1-4*, *STAT3*, *FANCF*. The CRISPR RNA sequences (shown in Supplementary Table 3) were synthesized by Eurofins Genomics and cloned to pLentiPuro plasmid (Addgene #52963) according to published instructions^1^. The plasmid identities were verified by Sanger sequencing.

The lentiviruses were prepared and transduced to HEK293T and BJ-5ta cells as described above. The cells were selected with 10 μg/ml puromycin (Thermo Fisher Scientific) or 10 μg/ml blasticidin (Sigma Aldrich) for 7 days prior to aliquoting and freezing.

### DNA repair protein library

The DNA repair protein Open Reading Frames (ORFs) and their corresponding sequences are found in the file “DNA repair domain library.xlsx”. In addition, the screen set contains 21 prokaryotic exonucleases as well as four engineered protein tags that are inert and function comparably to WTCas9. The corresponding full-lenght open reading frames (ORFs) were either picked from the human orfeome (hORF8.1) or synthesized by Genscript Inc. Since Cas9 is a large protein of >2000 amino acids (AAs), we expected the largest DNA repair proteins (>1000 AA) to lead to a failure in fusion protein expression. Therefore, the DNA repair proteins that exceeded the size of 600 AA were processed into smaller fragments (100-500AA) in between the domain boundaries. The fragments were synthesized by Genscript Inc.

### Expression plasmids

By utilizing the commercial pcDNA-DEST40 backbone (Invitrogen), we constructed a custom mammalian expression plasmid containing a WTCas9 sequence with N- and C-terminal nuclear localization signals, followed by a Gateway recombination site (referred as Cas9-GW plasmid). Genscript Inc. performed the insert synthesis and cloning to pcDNA-DEST40 backbone.

### High-throughput gateway cloning and plasmid preparation

The DNA repair protein inserts were cloned to Cas9-GW plasmids in 96-well plates in 5l reactions, with 1 μl Gateway LR clonase II enzyme mix (Invitrogen), 50 ng entry plasmid and 100 ng destination plasmids, respectively, and Tris-EDTA (TE) buffer (pH 8) added to a final volume of 5 μl. After overnight incubation, we transformed 1μl of the reaction mix to 15-20 μl of DH5-α competent cells (Invitrogen). Plasmids were extracted in 96-well format using the Wizard SV 96 plasmid miniprep kit (Promega) according to manufacturer’s instructions using the Biomek FXP liquid handling system (Beckman Coulter), and eluted to 100 μl TE buffer (pH 8). This yielded a destination plasmid concentration of ~150 ng in the majority of the wells.

### Cell transfections

HEK293T cells containing the GFP-BFP-sgRNA reporter cassette were split into 24-well plates at ~100 000 cells/well, 500 μl final volume. The next day, we replaced the media and prepared the transfection complexes in 96-well plates using Fugene HD (Promega) with the following modifications: with 25 μl Opti-MEM (Thermo Fisher), 2 μl Fugene HD (Promega), 400 ng plasmid and 12.5 pmol repair DNA (synthesized by Eurofins Genomics). After 20 min incubation, the cells were transfected in triplicate, with the end volume of 25 μl transfection mix in each well. Transfections were conducted with Biomek FXP liquid handling system (Beckman Coulter). The cells were grown for 5 days and GFP expression evaluated by FACS as described in the corresponding section.

### Fluorescence-activated cell sorting (FACS)

For the detection of corrected GFP, the Cas9-transfected cells were cultured for 5 days, trypsinized, resuspended in warm culture medium and immediately subjected to FACS. The data was acquired by CyAn II flow cytometer (Beckman Coulter) coupled with Hypercyte robotics (Intellicyt) or iQue screener plus (Intellicyt).

Well information was extracted from the plate fcs file using custom R code and Kaluza v.1.2 or 2.1 (Beckman Coulter). The number of GFP+ cells per well was obtained by batch gating. The gates used in analyzing the screen FACS data are shown on Fig. S8. The effect size for each Cas9 fusion was calculated by normalizing the GFP expression of the individual well either to the average GFP expression of all wells in the plate or to the overall experimental average. The statistical significance was calculated using standard one-way Anova test (Supplementary Table 5).

### Cas9WT fusion protein validation in HEK293T and RPE1 reporter cells

We chose 52 Cas9WT-DNA repair fusion constructs for validation. We picked the constructs from the original plates using Biomek i7 robotics and prepared new minipreps from the original constructs using the Wizard SV 96 Plasmid DNA Purification System (Promega), according to the manufacturer’s instructions. The constructs were Sanger sequenced to validate construct identity.

HEK293T cells containing the GFP-RFP-sgRNA reporter cassette were transfected as described in the corresponding section, with the following modifications: seeding on the 48 well plate, 30 000 cells/ well; transfection mix: 12.5 μl Opti-MEM (Thermo Fisher), 1.25 μl Fugene HD (Promega), 250 ng plasmid and 6 pmol repair DNA (synthesized by Eurofins Genomics). After 20 min incubation, the cells were transfected in four replicas. The transfections were performed in parallel plates to account for batch variation between cell culture plates. Transfections were conducted with Biomek i7 liquid handling system (Beckman Coulter). The cells were grown for 5 days and GFP expression evaluated by FACS.

RPE cells containing the GFP-RFP-sgRNA or GFP-BFP-sgRNA reporter cassette were electroporated with Lonza 4D 96-well electroporation system. 400 ng of the plasmid was pre-mixed with 40 pmol of the repair template on the 96 well PCR plate. RPE cells were trypsinized, washed with PBS and resuspended in the 1M electroporation buffer (5 mM KCl, 15 mM MgCl2, 120 mM Na2HPO4/NaH2PO4, pH 7.2, and 50 mM mannitol) to obtain a cell density of 200 000 cells per 20 μl of the buffer (ratio is for single-well reaction in the electroporation plate). 20 μl of the cell suspension was then dispensed in each well of the 96 well plate containing pre-mixed plasmids and repair template, gently mixed and transferred to the 96 well electroporation plate (Lonza). Electroporation was conducted using the pulse code EA-104. Instantly after electroporation, 80 μl of pre-warmed medium (DMEM, supplemented with 10% FBS, no antibiotics) was added into each well. Cells were collected and plated on 48-well plates (Costar) containing 300 μl of pre-warmed medium (DMEM, supplemented with 10% FBS, no antibiotics). Liquid handling steps were done with the Biomek i7 platform. Electroporated cells were cultured for 5 days, after which the GFP expression was evaluated by FACS.

### Cas9WT fusion protein validation in endogenous loci (HEK293T cells)

We chose 31 Cas9WT-DNA repair fusion constructs for validation in endogenous loci in HEK293T cells. We used four different HEK293T lines, each stably expressing one of the sgRNAs targeting *ELANE*, *CTCF1*, *Ench4-1*, *RNF2* genomic loci (CRISPR RNA sequences in Supplementary Table 3). Cells seeded to 24-well plates at ~100 000 cells/well were transfected as described above, with the following modifications: 25 μl Opti-MEM (Thermo Fisher), 2.5 μl Fugene HD (Promega), 480 ng of Cas9 fusion plasmid and 10 pmol repair DNA. After 20 min incubation, the cells were transfected in triplicate. After 5 days of incubation, DNA was extracted with PureLink 96 Genomic DNA kit (Invitrogen) and analyzed using droplet digital PCR.

### Droplet digital PCR

We performed droplet digital PCR (ddPCR) assays to estimate non-homologous end-joining (NHEJ) and homology directed repair (HDR) efficiencies at the *RNF2*, *Enh4-1*, *FANCF*, *ELANE*, *STAT3*, *CTCF1* and *GFP* guide target loci. The assay schematic is shown in Fig. S9. The ddPCR was performed on the QX200 system (BioRad Laboratories). Final reaction mixture volume was 20 μL: 10 μL of 2× ddPCR Super Mix for Probe (BioRad Laboratories), 8 μl of DNA (concentration normalized to 8 ng/μl), primers (900 nM), reference probe (250 nM) and HDR or NHEJ probe (250 nM). Each reaction was then loaded into a sample well of an eight-well disposable cartridge (DG8^™^; Bio-Rad Laboratories) along with 70 μl of droplet generation oil (Bio-Rad Laboratories). Droplets were formed using a QX200^™^ Droplet Generator (Bio-Rad Laboratories). Droplets were transferred to a 96-well PCR plate, heat-sealed with foil and amplified using a conventional thermal cycler. The thermocycling protocol was the following: 1) 95°C - 10 min, 2) 94°C - 30 sec, 56°C - 3min, step repeated 42 times 3)98°C - 10 minutes 4) 4°C-hold. The resulting PCR products were loaded on a QX200 Droplet Reader (Bio-Rad Laboratories), and the data analyzed using QuantaSoft^™^ software (Bio-Rad Laboratories). Primer and probe sequences are listed in Supplementary Table 4. The data was analyzed with Quantasoft software. Representative gatings are shown in Fig. S10.

### In vitro mRNA transcription

Cas9WT fusion mRNA was prepared with HiScribe^™^ T7 ARCA mRNA Kit with tailing (NEB-bionordika), according to the manufacturer’s instructions. Total of 8000 ng stock plasmid was digested with 2 μl FastDigest MssI enzyme in the supplemented restriction-digestion buffer, total reaction volume 20 μl. Incubation was carried out at 37°C overnight. Length of the digested product was confirmed by gel electrophoresis. For the IVT reaction, 1000 ng of the linearised plasmid was mixed with 10 μl of 2xARCA/NTP mix and 2 μl of T7 RNA Polymerase mix, and the reaction was incubated for 30 min at 37°C. Sequentially, 2 μl of DNAse enzyme was added and the mixture was incubated at 37°C for 15 min. Poly(A) tailing step was performed by adding 20 μl of milliQ (RNAse free), 5 μl of 10x PolyA polymerase reaction buffer and 5 μl of 10x PolyA polymerase directly to the IVT reaction, which was incubation at 37°C for 30 min. mRNA was purified using LiCl solution, as described in the manufacturer’s protocol. Aliquots were frozen in −80°C for future use.

### Cas9WT fusion protein validation in endogenous loci (BJ-5ta cells)

We chose 7 Cas9WT fusion constructs for validation in endogenous loci in immortalized BJ-5ta fibroblasts. We used five different BJ5ta lines, each stably expressing one of the sgRNAs (IDT) targeting *FANCF*, *Enh4-1*, *STAT3*, *ELANE*, or *RNF2* genomic loci (CRISPR RNA sequences in Supplementary Table 4). The Cas9 fusion constructs were transfected as mRNA; IVT was performed as described above. BJ-5ta-sgRNA cell lines were electroporated with Lonza 4D 96-well electroporation system as described in corresponding section, with following modifications: 1000 ng of mRNA and 100 pmol of the corresponding HDR repair template per well, cell density 500 000 of cells per 20 μl of the electroporation buffer. Cells were pulsed using code CA-137. Cell culture was carried out on 6-well plates for 5 days, followed by the DNA isolation (DNeasy Blood & Tissue Kit (250), Qiagen). Editing efficiency was evaluated by ddPCR as described in the corresponding section.

### Cas9-POLD3 validation in peripheral blood mononuclear cells

Peripheral blood mononuclear cells (PBMCs) were isolated from whole blood by density gradient separation method using Lymphoprep^™^ (StemCell Technologies, #07851). Obtained cells were cultured in the T cell expansion media (RPMI 1640 Medium, GlutaMAX Supplement, HEPES (Gibco, #72400- 021) supplemented with 10% FBS (Fetal Bovine Serum, certified, heat inactivated, Gibco, #10082147), 25 μl/ml anti-human CD3/CD28 (StemCell Technologies, #10970), 200 U/ml IL-2 (Recombinant Human IL-2, Peprotech, # 200-02), 5 ng/ml IL-7 (Recombinant Human IL-7, Peprotech, #200-07), 5 ng/ml IL-15 (Recombinant Human IL-15, Peprotech, #200-15) with 1× P/S (Penicillin-Streptomycin-Glutamine 100X, Gibco #10378016) for 3 days prior the electroporation at 1×10^6^ cells/ml density.

Electroporation was performed using the Lonza 4D Nucleofector System, Core Unit (Lonza cat. no AAF-1002B) with a 96-well Shuttle Device (Lonza cat. no AAM-1001S). 1×10^6^ of cells were resuspended in 20 μl of the home-made electroporation buffer (5 mM KCl, 15 mM MgCl2, 120 mM Na2HPO4/NaH2PO4, and 50 mM mannitol, pH 7.2), combined with 1000 ng of nuclease/fusion encoding mRNA, 100 pmol of the corresponding sgRNA and 100 pmol of the repair template (ssODN) and electroporated using EO-115 pulse code. After the electroporation, 80 μl of the recovery media (RPMI 1640 Medium, GlutaMAX Supplement, HEPES (Gibco, #72400-021) supplemented with 10% FBS (Fetal Bovine Serum, certified, heat inactivated, Gibco, #10082147), IL-2 at 500 U/ml, without any antibiotics) was added to each well and cells were incubated for 15 min in the 37°C, 5% CO2. Subsequently, cell suspension from each well was transferred to the 400 μl of the recovery medium pre-dispensed on the 24 well plates. Half of the medium volume from each well was replaced with fresh recovery medium supplemented with 1× P/S (Penicillin-Streptomycin-Glutamine 100X, Gibco #10378016) daily, until the sample collection time point at 96h post-electroporation. DNA extraction was done using the DNeasy 96 Blood & Tissue Kit (4) (Qiagen, #69581) according to the kit instructions.

### Cas9-POLD3 validation in human embryonic stem cells

The human male embryonic stem cell (hESC) H13 line^2^ was obtained from the WiCell Research Institute (Madison, Wisconsin) and cultured on mitomycin-C treated mouse embryonic fibroblasts (MEFs). HESCs were maintained in DMEM/F-12 (Sigma, #D8900-50L) media containing 15% FBS (Capricorn Scientific, #FBS-ES-HI12A), 5% KnockOut Serum Replacement (Gibco, #10828-028), 1% MEM Non-essential Amino Acids (Gibco, #11140-068), 1% GlutaMAX (Gibco, #35050-038), 1% antibiotic-antimycotic solution (Gibco, #15240-062), 55 μM 2-mercaptoethanol (Gibco, #31350-010), and 5 ng/ml of recombinant human bFGF (Miltenyi Biotec, #130093842). The cells were first passaged 5-6 days prior to the start of the experiment using collagenase type IV (1 mg/ml, Life Technologies, # 17104019) and transferred into MEF-coated 6-well plates. The media was exchanged 24h prior the electroporation and supplemented with 10 μM Rho Kinase (ROCK)-inhibitor (ATCC, Y-27632).

Immediately before the electroporation, hESC colonies were detached from plates using 1 mg/ml collagenase IV (Gibco, #17104019) and sedimented in 10 ml of PBS, pH 7,4, for 5 minutes. After brief centrifugation (100xg, 1 min) PBS was removed and hESC were disrupted into single cells using 0.25 % trypsin/ EDTA solution (Gibco, 25200-114) for 3 min at the 37°C, 5% CO2. Trypsin was neutralized with hESC media and the cells were pelleted at 150xg for 5 minutes. The cell pellet was resuspended in 2 ml PBS, pH 7.4, and cells were counted using Trypan blue on the automated cell counter (Countess). Electroporation was performed using the MaxCyte Expert electroporation system with OC-25×3 Electroporation Cuvettes (MaxCyte, #238537). For each reaction, 0.4×10^6^ of cells were resuspended in 25 μl of the MaxCyte electroporation buffer (MaxCyte, #EPB1), combined with 1000 ng of Cas9 mRNA, 100 pmol sgRNA (IDT) targeting the RNF2 locus and 100 pmol of the repair DNA template and electroporated using Optimization-8 pulse code. After the electroporation, cells in the electroporation cuvettes were incubated for 15 min in the 37°C, 5% CO2. Then, the cell suspension was transferred to 24-well plates pre-coated with Matrigel (Corning, #734-1440) and supplemented with 1 ml hESC culture media (no antibiotics, supplemented with 10 μM ROCK-inhibitor). The media was exchanged 24 and 72 h after the electroporation. The DNA was extracted after 96h using the DNeasy Blood & Tissue Kit (Qiagen, # 69504) according to the manufacturer’s instructions.

### Recombinant protein production

The hit Cas9 fusion proteins were synthesized by GeneArt Inc. and cloned to pET301/CT-DEST vector (Invitrogen) or pTH21 vector (PMID 16150509)^3^ for *E.coli* expression. The Cas9 sequence as well as the sequence and position of nuclear localization signal and GS linker were similar to the Cas9-GW sequence presented in Appendix I. The exact sequences are listed in the file DNA repair protein domain library.xlsx.

All proteins contained C-terminal His-tags and were expressed in *E. coli* BL21 (DE3) T1R pRARE2 at 18 °C and purified by the Protein Science Facility (PSF) at Karolinska Institutet, Stockholm. Purification was performed using the HisTrap HP column (GE Healthcare) followed by gel filtration step with a HiLoad 16/60 Superdex 200 (GE Healthcare). Purity of the protein preparation was examined using SDS-PAGE followed by Commassie staining. All purified proteins were concentrated, aliquoted and stored in a storage buffer (20 mM HEPES, 300 mM NaCl, 10% glycerol, 2 mM TCEP, pH 7.5) at −80°C.

### RNP complex preparation & transfection

We obtained the mGFP-targeting CRISPR and TRACR RNAs from Integrated DNA Technologies (IDT). The CRISPR/TRACR hybridization was performed by incubation of crRNA tracrRNA in equimolar concentrations at 95°C followed by gradual cooling down to RT. For ribonucleoprotein (RNP) complex formation nuclease (protein or fusion) was mixed gRNA in 1:2 molar ratio and incubated 15 min RT. For transfection, reporter HEK293T and RPE1 cells were plated in 48-well plates at 100 000 cells/well and reverse transfected with 13 pmol of RNP and 6 pmol of repair DNA template using the CRISPRmax transfection reagent (Thermo Scientific) according to the manufacturer’s instructions. For the detection of corrected GFP, the transfected cells were cultured for 4 days, trypsinized, resuspended in warm culture medium and immediately subjected to FACS. The data was acquired by CyAn II flow cytometer (Beckman Coulter) coupled with Hypercyte robotics (Intellicyt) and analyzed with Kaluza v.1.2 (Beckman Coulter).

### Functional comparison between Cas9-POLD3 and previously published fusions

HEK293T cells containing the GFP-BFP-sgRNA reporter cassette were transfected as described in the corresponding section, with the following modifications: seeding on the 48 well plate: 30 000 cells/ well; transfection mix: 12.5 μl Opti-MEM (Thermo Fisher), 1.25 μl Fugene HD (Promega), 250 ng plasmid and 6 pmol repair DNA (synthesized by Eurofins Genomics). After 15 min incubation, the cells were transfected in four replicas. The transfections were performed in parallel plates to account for batch variation between cell culture plates.The cells were grown for 5 days and GFP expression evaluated by FACS.

### DNA breakpoint quantification

We used immortalized fibroblasts (BJ-5ta) expressing a guide that targets the CTCF1 locus. The cells were electroporated as described above with a pool of sgRNAs (IDT) targeting the Enh 4-1, STAT3 and RNF2 loci, 100 pmol RNA per reaction. The purpose of using the pool is to increase the total number of edited alleles per nucleus for better visualization. After the electroporation, cells were plated on 6-well plates containing clean glass coverslips and cultured as described above.

Samples were collected for staining at 8h, 12h, 24h, 48h, 72h and 96h post-electroporation. We used the coverslips for γH2AX foci quantification, and the cells in the surrounding space in the well for NHEJ quantification by ddPCR (described in the corresponding section). For confocal microscopy, each sample slide was washed twice with PBS and fixed with 4% PFA for 15 min at room temperature (RT), followed by another PBS washing step and storage at +4°C in PBS, pH 7,4 until the sample collection was finished. The samples were then permeabilized (0.1% Triton X-100 in PBS for 45 min RT) and blocked (PBS with 0,1% Triton X-100, 5% BSA and 5% goat serum for 45 min RT). After blocking, Anti-phospho-H2AX (Ser139) (Millipore, 05-636-25UG) was added in 1:500 ratio, diluted in 0,1% Tween-20, 0.5% BSA and 0.5% goat serum in PBS and and the samples were incubated overnight at 4°C. The following day, coverslips with samples were washed three times with 0,1% Triton X-100 in PBS with shaking and Goat anti-Mouse IgG (H+L) Cross-Adsorbed Secondary Antibody, Alexa Fluor 488 (Thermo Fisher, A-11001) was added in 1:1000 dilution in 0.1% Triton X-100 in PBS and incubated for 1h at RT, protected from light. After three additional washes in PBS, nuclei were stained with DAPI (1 μg/ml in PBS). The slides were mounted using Fluoromount^™^ Aqueous Mounting Medium (Sigma Aldrich, F4680).

Widefield fluorescence imaging (Z-stacks) was done using Zeiss AxioObserver Z1, with a Plan Apo 40X/1.4 Oil objective and a Hamamatsu ImagEMX2 EMCCD. Imaging settings (filter set, lamp power and camera exposure) were kept constant for image collection for all samples. Image analysis was done with batch mode in FIJI (ImageJ) with two main steps: 1. Segmentation of Region of Interest (ROI) to outline cell nuclei with DAPI channel with the FIJI plugin Auto Threshold(Mean) and watershed algorithm; 2. Counting the γH2AX foci in ROIs identified in Step 1 with the FIJI plugin Find Maxima (prominence=2000). Average foci/nucleus ratio was calculated for each sample based on the quantified results.

### Cas9WT fusion plasmid titration in RPE1 reporter cells

The Cas9 fusions that showed a trend for improving over Cas9 WT were Sanger sequenced to validate construct identity. We then prepared new minipreps from the original constructs using Qiaprep spin Miniprep kit (Qiagen) according to manufacturer’s instructions.

For the time-course experiment, RPE-GFP-BFP-gRNA cells were electroporated using the Lonza 4D 96-well electroporation system as described in the corresponding section with the following modifications: 200 000 cells per each well in 24 well plates; 400 ng of plasmid, 100 pmol of GFP repair template; pulse code changed to EA-104. Cells were gradually collected at 24h, 48h, 72h post-electroporation. DNA extracted using DNeasy Blood & Tissue Kit (250), (Qiagen, # 69504). Editing efficiency was evaluated by ddPCR as described in the corresponding section.

For the gradient titration, RPE-GFP-BFP-gRNA cells were electroporated with Lonza 4D 16-well electroporation system as described in corresponding section with the following modifications: 200 000 cells per each well in 24 well plates; 15, 30, 60, 120 pmol of plasmid, 40 pmol of GFP repair template; pulse code changed to EA-104. FACS analysis was performed after 5 days by iQue screener plus (Intellicyt) and analysed using Kaluza v 2.1 software (Beckman Coulter), as described in corresponding section.

### Affinity-purification mass spectrometry and BioID proximity labelling

The hit Cas9 fusion proteins were synthesized and cloned to pDNOR221 Gateway entry vector (Invitrogen) by GeneArt Inc. Sequences are listed in DNA_repair_domain_library.xlsx. The inserts were cloned to MAC-Tag-C vector^4^ (PMID 29568061, Addgene ID #108077), which adds a c-terminal MAC tag (contains Strep-tag and modified minimal biotin ligase) to the Cas9 fusion.

For generation of the stable cell lines inducibly expressing the MAC-tagged versions of the baits, Flp-In^™^ T-REx^™^ 293 cell lines (Invitrogen, Life Technologies, R78007) were first transduced with the lentivirus containing the GFP-BFP-sgRNA cassette, as described in corresponding section. The cells were co-transfected with the MAC-tagged expression vector and the pOG44 vector (Invitrogen) using the Fugene HD transfection reagent (Promega). Two days after transfection, cells were selected in 50 μg/ml streptomycin and hygromycin (100 μg/ml) for 2 weeks.

We tested the presence of the fusion and the functionality of the GFP-BFP-sgRNA reporter cassette by FACS prior to the experiment. The Cas9 expression was induced with 1 μg/ml tetracycline for 24h, followed by GFP repair template transfection by RNAiMax transfection reagent (Thermo Fisher Scientific) according to manufacturer’s instructions.

For AP-MS experiments, each stable cell line was expanded to 80% confluence in 20 × 150 mm cell culture plates. 2×5 plates were used for AP-MS approach, in which 1 μg/ml tetracycline was added for 30h induction, and 2×5 plates for BiolD approach, in which in addition to tetracycline, 50 μM biotin was added for 30 h before harvesting. Cells from 5 × 150 mm fully confluent dishes (~5 × 10^7^ cells) were pelleted as one biological sample. Thus, each bait protein has two biological replicates in two different experiments. All samples from one experiment were processed in parallel. Samples were snap frozen and stored at −80 °C. Analyses of the baits were performed as two biological replicates.

For the Affinity Purification Mass Spectrometry (AP-MS) approach, the sample was lysed in 3 ml of lysis buffer 1 (0.5% IGEPAL, 50 mM Hepes, pH 8.0, 150 mM NaCl, 50 mM NaF, 1.5 mM NaVO3, 5 mM EDTA, supplemented with 0.5 mM PMSF and protease inhibitors; Sigma).

For BiolD approach, Cell pellet was thawed in 3 ml ice-cold lysis buffer 2 (0.5% IGEPAL, 50 mM Hepes, pH 8.0, 150 mM NaCl, 50 mM NaF, 1.5 mM NaVO3, 5 mM EDTA, 0.1% SDS, supplemented with 0.5 mM PMSF and protease inhibitors; Sigma). Lysates were sonicated and treated with benzonase.

Cleared lysate was obtained by centrifugation and loaded consecutively on spin columns (Bio-Rad) containing lysis buffer 1, prewashed 200 μl Strep-Tactin beads (IBA, GmbH). The beads were then washed 3 × 1 ml with lysis buffer 1 and 4 × 1 ml with wash buffer (50 mM Tris-HCl, pH 8.0, 150 mM NaCl, 50 mM NaF, 5 mM EDTA). Following the final wash, beads were then resuspended in 2× 300 μl elution buffer (50 mM Tris-HCl, pH 8.0, 150 mM NaCl, 50 mM NaF, 5 mM EDTA, 0.5 mM Biotin) for 5 mins and eluates collected into Eppendorf tubes, followed by a reduction of the cysteine bonds with 5 mM Tris(2-carboxyethyl)phosphine (TCEP) for 30 mins at 37°C and alkylation with 10 mM iodoacetamide. The proteins were then digested to peptides with sequencing grade modified trypsin (Promega, V5113) at 37°C overnight. After quenching with 10% TFA, the samples were desalted by C18 reversed-phase spin columns according to the manufacturer’s instructions (Harvard Apparatus). The eluted peptide sample was dried in vacuum centrifuge and reconstituted to a final volume of 30 μl in 0.1% TFA and 1% CH3CN.

Analysis was performed on a Q-Exactive mass spectrometer using Xcalibur version 3.0.63 coupled with an EASY-nLC 1000 system via an electrospray ionization sprayer (Thermo Fisher Scientific). In detail, peptides were eluted and separated with a C18 precolumn (Acclaim PepMap 100, 75 μm x 2 cm, 3 μm, 100 Å, Thermo Scientific) and analytical column (Acclaim PepMap RSLC, 75 μm × 15 cm, 2 μm, 100 Å; Thermo Scientific), using a 60 min buffer gradient ranging from 5 to 35% buffer B, followed by a 5 min gradient from 35 to 80% buffer B and 10 min gradient from 80 to 100% buffer B at a flow rate of 300 nl min-1 (buffer A: 0.1% formic acid in 98% HPLC grade water and 2% acetonitrile; buffer B: 0.1% formic acid in 98% acetonitrile and 2% water). For direct LC-MS analysis, 4 μl peptide samples were automatically loaded from an enclosed cooled autosampler. Data-dependent FTMS acquisition was in positive ion mode for 80 min. A full scan (200–2000 m/z) was performed with a resolution of 70,000 followed by top10 CID-MS2 ion trap scans with resolution of 17,500. Dynamic exclusion was set for 30 s. Acquired MS2 spectral data files (Thermo.RAW) were searched with Proteome Discoverer 1.4 (Thermo Scientific) using SEQUEST search engine of the selected human component of UniProtKB/SwissProt database (http://www.uniprot.org/, version 2015–09). The following parameters were applied: Trypsin was selected as the enzyme and a maximum of 2 missed cleavages were permitted, precursor mass tolerance at ±15 ppm and fragment mass tolerance at 0.05 Da. Carbamidomethylation of cysteine was defined as a static modification. Oxidation of methionine and biotinylation of lysine and N-termini were set as variable modifications. All reported data were based on high-confidence peptides assigned in Proteome Discoverer with FDR<1%.

The mass spectrometric data was searched with Proteome Discover 1.4 (Thermo Scientific) using the SEQUEST search engine against the UniProtKB/SwissProt human proteome (http://www.uniprot.org/, version 2015-09). Search parameters were set either as in PMID: 29568061(QE runs; BioID)^4^ or in PMID: 28054750 (Orbitrap Elite runs; AP-MS)^5^. All data was filtered to medium-(AP-MS) or high-confidence (BioID) peptides according to Proteome Discoverer FDR 5% or 1%, respectively. The lists of identified proteins were conventionally filtered to remove proteins that were recognized with less than two peptides and two PSMs. The high-confidence interactors were identified using SAINT and CRAPome as in Liu et al., 2018. Each sampl22e’s abundance was normalized to its bait abundance. These bait-normalized values were used for data comparison and visualization.

### GUIDE-seq

HEK293T cells were cultured in DMEM (Life Technologies) supplemented with 10% FBS, and penicillin/streptomycin at 37°C with 5% CO2. For each replicate, 300 000 cells were transfected in 20 μl Solution P3 (Lonza) on a Lonza Nucleofector 4-D using the program CM-137, according to the manufacturer’s instructions. 400 ng of pcDNA-pDEST-Cas9wt-POLD3, 150 ng of gRNA encoding plasmid (pU6-gRNA-HEK site 4) and 5 pmol of dsODN were transfected.

The blunt-ended dsODN used in our GUIDE-seq experiments was the same as which was used in the original publication^6^, and then was prepared by annealing the two modified oligonucleotides of the following compositions:

5’-P-G*T*TTAATTGAGTTGTCATATGTTAATAACGGT*A*T −3’

and

5’- P-A*T*ACCGTTATTAACATATGACAACTCAATTAA*A*C −3’

P represents a 5’ phosphorylation and * indicates a phosphorothioate linkage.

Genomic DNA was isolated using Qiagen DNeasy Blood & Tissue Kit (250) (#69506) and sheared with a Bioruptor Pico instrument (15 sec ON, 90 sec OFF, 7 cycles) to an average length of around 400-500 bp. The sheared gDNA were then end-repaired, A-tailed and ligated to half-functional adapters, incorporating an 8-nt random molecular index according to the details of the GUIDE-seq protocol provided in the original GUIDE-seq paper^6^. Next, two rounds of nested anchored PCR, with primers complementary to the oligo tag, were used for target enrichment, as described in original publication^6^. Next, the samples were QC by BioAnalyzer 2100, DNA concentrations quantified by Qubit and samples were pooled together and sequenced using the Illumina MiSeq instrument.

Data analysis was performed using the GUIDEseq analysis pipeline^7^. We used custom scripts to extract UMI from the demultiplexed fastq reads and trimmed adapters with cutadapt v2.8 (TTGAGTTGTCATATGTTAATAACGGTAT and ACATATGACAACTCAATTAAAC). Afterwards, the data was aligned to the human genome (hg38) using bowtie2 v2.3.5.1 (with options –local –very-sensitive-local). The GUIDEseq v1.18 package from Bioconductor was used to call off-targets with the default settings. Final off-targets were normalized against control data (HEK293T cells transfected with dsODN only). Control data was processed in the same pipeline as the modified Cas9 samples.

### Amplicon sequencing

BJ-5ta cell lines expressing guides against RNF2 and ELANE loci were electroporated as described earlier. The DNA was extracted using the DNeasy Blood Tissue Kit (Qiagen, # 69504) according to the kit instructions.

Library preparation was performed using a two-step PCR method. For the first PCR reaction, a pair of target-specific primers was designed to amplify the 150bp area surrounding the cutting site. Each target primer additionally includes an extension at the 5’ end: for forward primers, this contains the Illumina Read 1 primer sequence (see below, nucleotides in bold) and an 8bp Unique Molecular Identifier (UMI, nucleotides underlined), and for the reverse primers, this contains the Illumina Read 2 primer sequence only (see below, nucleotides in bold):

RNF2 fwd 5’-3’: **ACA CTC TTT CCC TAC ACG ACG CTC TTC CGA TCT** <u>NNN NNN NN</u>G ACA AAC GGA ACT CAA CCA T
RNF2 rev 5’-3’: **GTG ACT GGA GTT CAG ACG TGT GCT CTT CCG ATC T**TG TTC TAT TTA AGT TTT CAT GTT CT
ELANE fwd 5’-3’: **ACA CTC TTT CCC TAC ACG ACG CTC TTC CGA TCT** <u>NNN NNN NN</u>C TCC CCG GCA GAA ACG TC
ELANE rev 5’-3’: **GTG ACT GGA GTT CAG ACG TGT GCT CTT CCG ATC T**GA GAA TCA CGA TGT CGT TGA GC

Each PCR reaction was composed as follows:

| Component | Final conc. | µl per 1 reaction |
| --- | --- | --- |
| H<sub>2</sub>O (milliQ) |  | Add to obtain total of 50µl |
| 5X Phusion GC Buffer (Thermo Fisher Scientific, #00928987) | 1X | 10 |
| 10mM dNTPs (Thermo Fisher Scientific, #R0192) | 200 µM | 1 |
| 10µM Primer fwd | 0.5 µM | 2.5 |
| 10µM Primer rev | 0.5 µM | 2.5 |
| Betaine (Sigma Aldrich, #B0300) | 1M | 10 |
| Phusion Hot Start II DNA Polymerase (2 U/µL) (Thermo Fisher Scientific, #00928987) | 0.02 U/µL | 0.5 |
| Template DNA | 500 ng |  |

Thermo-cycling conditions:

| Cycle step | Temp. | Time | Cycles |
| --- | --- | --- | --- |
| 1. Initial denaturation | 98 °C | 30 s | 1 |
| 2. Denaturation | 98 °C | 10 s | 30 |
| 3. Annealing | 57 °C | 10 s |  |
| 4. Extension | 72 °C | 20 s |  |
| 5. Final extension | 72 °C | 5 min | 1 |
| 6. Hold | 4 °C | forever |  |

For the second PCR reaction, the amplified products were purified using AMPure XP magnetic (Beckman Coulter, #A63882) according to the manufacturer instructions, pooled and annealed with i5 and i7 Illumina Index primers. Both primers contain flow-cell binding region (see below, highlighted in bold), index region (see below, underlined) and Illumina Read1 or Read2 primer binding regions, correspondingly, (see below, italics):

i5-PCR Index 11:
**AAT GAT ACG GCG ACC ACC GAG ATC TA**<u>A CCT GGT T</u>*AC ACT CTT TCC CTA CAC GAC GCT CTT CCG ATC*T*
i5-PCR Index 13:
**AAT GAT ACG GCG ACC ACC GAG ATC TA**<u>C GGA ACA A</u>*AC ACT CTT TCC CTA CAC GAC GCT CTT CCG ATC*T*
i7-PCR Index 13:
**CAA GCA GAA GAC GGC ATA CGA GAT** <u>TTC CTC CT</u>*G TGA CTG GAG TTC AGA CGT GTG CTC TTC CGA TC*T*
i7-PCR Index 14:
**CAA GCA GAA GAC GGC ATA CGA GAT** <u>TGC TTG CT</u>*G TGA CTG GAG TTC AGA CGT GTG CTC TTC CGA TC*T*
i7-PCR Index 15:
**CAA GCA GAA GAC GGC ATA CGA GAT** GGT GAT GA*G TGA CTG GAG TTC AGA CGT GTG CTC TTC CGA TC*T*
i7-PCR Index 16:
**CAA GCA GAA GAC GGC ATA CGA GAT** <u>AAC CTA CG</u>*G TGA CTG GAG TTC AGA CGT GTG CTC TTC CGA TC*T*

Each PCR reaction was composed as following:

| Component | Final conc. | µl per 1 reaction |
| --- | --- | --- |
| H<sub>2</sub>O (milliQ) |  | Add to obtain total of 50µl |
| 5X Phusion GC Buffer (Thermo Fisher Scientific, #00928987) | 1X | 10 |
| 10mM dNTPs (Thermo Fisher Scientific, #R0192) | 200 µM | 1 |
| 10µM Primer i5 | 0.4 µM | 2 |
| 10µM Primer i7 | 0.4 µM | 2 |
| Betaine (Sigma Aldrich, #B0300) | 1M | 10 |
| Phusion Hot Start II DNA Polymerase (2 U/μL)<br>(Thermo Fisher Scientific, #00928987) | 0.02 U/μL | 0.5 |
| Template DNA | 500 ng |  |

Thermo-cycling conditions:

| Cycle step | Temp. | Time | Cycles |
| --- | --- | --- | --- |
| 1. Initial denaturation | 98 °C | 30 s | 1 |
| 2. Denaturation | 98 °C | 10 s | 10 |
| 3. Annealing | 58 °C | 10 s |  |
| 4. Extension | 72 °C | 20 s |  |
| 5. Final extension | 72 °C | 5 min | 1 |
| 6. Hold | 4 °C | forever |  |

PCR products were purified using AMPure XP magnetic beads (Beckman Coulter, #A63882) according to the manufacturer’s instructions and DNA concentrations were measured with Qubit HS kit (Thermo Fisher Scientific, #2113695). Final products were pooled in equimolar concentrations into a single library with final concentration 250 nM (50x concentrate). Sequencing was done using an Illumina MiSeq v2 Micro flow cell, including 10% PhiX. The data analysis was performed using the AmpliCan software package^8^.

### Statistics

#### High-throughput screening

##### This article describes one high-throughput screen

1) A screen with ~450 DNA repair proteins fused with Cas9 and tested in clonal HEK293T cells containing the GFP-BFP-sgRNA. The screen contained three biological replicates (=parallel cell culture wells in different culture plates) and was conducted twice (n = 2×3) (Fig. 1b).

To calculate statistical significance, we first calculated the average of GFP+ cells in each biological replica. The data points were then compared to the combined mean of GFP+ cells in the whole screen (“experiment average”) or the 48-well cell culture plate (“plate average”), as indicated in Fig. 1B. We compared the mean of each replicate group to the combined mean of all other replicate groups using one-way Anova testing. The statistical parameters for these experiments are shown in Supplementary Table 5. Note that in this case, unlike pairwise comparisons in usual Anova with multiple treatments, there is no multiple comparison problem since we compare in-turn the mean of each replicate group to a very large background group. Hence, each test statistic is dominated by the smaller replicate group so that the test samples are essentially disjoint.

For visualization, the data points were normalized to the average of GFP+ cells in the whole screen (“experiment average”) or the 48-well cell culture plate average (“plate average”), as indicated in Fig. 1b.

##### Validation

Fig. 1c-d: We chose 52 Cas9WT-DNA repair protein fusions that performed well in the main screen, and tested them in clonal HEK293T cells containing the GFP-BFP-sgRNA, with each tested fusion having 4 replicates (parallel cell culture wells in different 48-well plates). In addition, we tested the fusions in clonal RPE-GFP-RFP-sgRNA cells once (n=1×2) and in clonal RPE-GFP-BFP-sgRNA cells once (n=1×2). We used different clonal lines than that in the major screen to account for clone-specific phenomena.

Fig. 1f: We chose 31 Cas9WT-DNA repair protein fusions that performed well in the main screen and in the first validation, and tested them in pooled HEK293T cells containing guides against the endogenous loci (*ELANE*, *RNF2*, *Enh 4-1*, *CTCF1*). Each tested fusion had 3 replicates (parallel cell culture wells in different 48-well plates) tested in four independent experiments (one independent experiment for each gene locus). For both validation experiments, the statistics were calculated similarly to the main screens.

Fig. 2b: We chose 7 Cas9WT-DNA repair protein fusions that performed well in the previous validation experiments, and tested them in BJ-5ta-sgRNA cells containing guides against the endogenous loci (*ELANE*, *RNF2*, *Enh 4-1*, *FANCF*, *STAT3*). Each tested fusion had 4 replicates, except STAT3 where n=3, (parallel cell culture wells in different 6-well plates) tested in 1 independent experiment (one independent experiment for each gene locus).

Fig. 3b: Four replicates (=parallel wells in a cell culture plate) for each condition (n=1×4), one independent experiment. P-values denote significance of the total editing (HDR+NHEJ) increment between the Cas9WT and other fusions (for 24h to 48h period). Statistical values derived using one-way ANOVA test.

Fig. 3e: Five fluorescent microscopy images were taken from each individual glass slide (n=5), each glass slide corresponds to one particular time point for each tested condition; data from one independent experiment. Statistical significance of the difference between Cas9WT and Cas9-POLD3 is calculated using ANOVA test for the equality of the means at a particular time point.

Fig. 3f-i: One replicate (=well in a cell culture plate) for each individual time point and electroporation cargo (n=1), ddPCR reactions individually conducted for each gene locus; data from one independent experiment (same set as shown in Fig. 3e).

Fig. 4a-b: The mass spectrometry experiments were conducted in biological duplicates (=parallel cell culture dishes). The data analysis is explained in the corresponding section.

Fig. 5a-b: One replicate (=one well in a cell culture plate) for each condition (n=1), one independent experiment.

Fig. 5c-f: One replicate (=one well in a cell culture plate) for each condition (n=1), one independent experiment.

Fig. 6a: Validation of Cas9-POLD3 against the panel of various HDR improving fusions in reporter HEK293T cells. n=4, one of two independent experiments, bar denotes mean value, error bars represent ± S.D. Statistical significance is calculated with unpaired, two-sided Student’s t-test.

Fig. 6b: Three Cas9 fusion proteins were tested in RPE-1 and HEK293T cells. n=5, one of two independent experiments, bar denotes mean value, error bars represent ± S.D. Statistical significance is calculated with Anova that is adjusted for multiple comparisons.

Summary of all statistical data is available in Supplementary Table 5.

Data availability statement: The raw data can be obtained from the corresponding author upon request. Accession codes will be available before publication.

