## Supplementary Material for "Rapid genome editing by CRISPR-Cas9-POLD3 fusion"

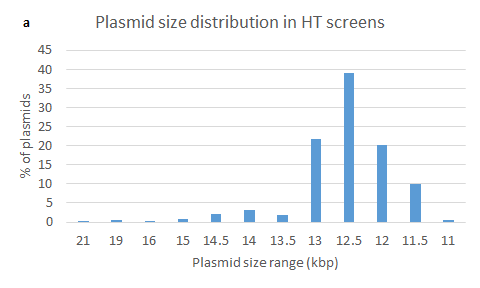

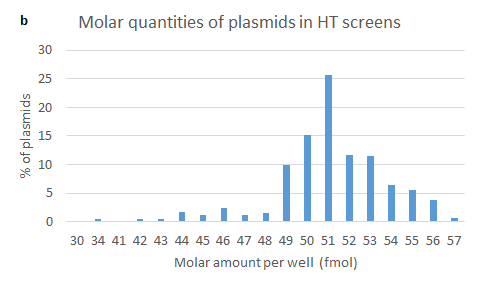
 **Figure S1. Plasmid sizes and molar quantities of plasmids used in the high throughput screens.
a.** Percentage distribution of plasmids used in the high throughput screens according to their size; 90% of plasmids range from 11.5 to 13 kbp in size. **b.** Percentage distribution of plasmids used in the high throughput screens according to their molar amounts per well; 90% of plasmids were used in quantities ranging from 49 fmol to 56 fmol per well.

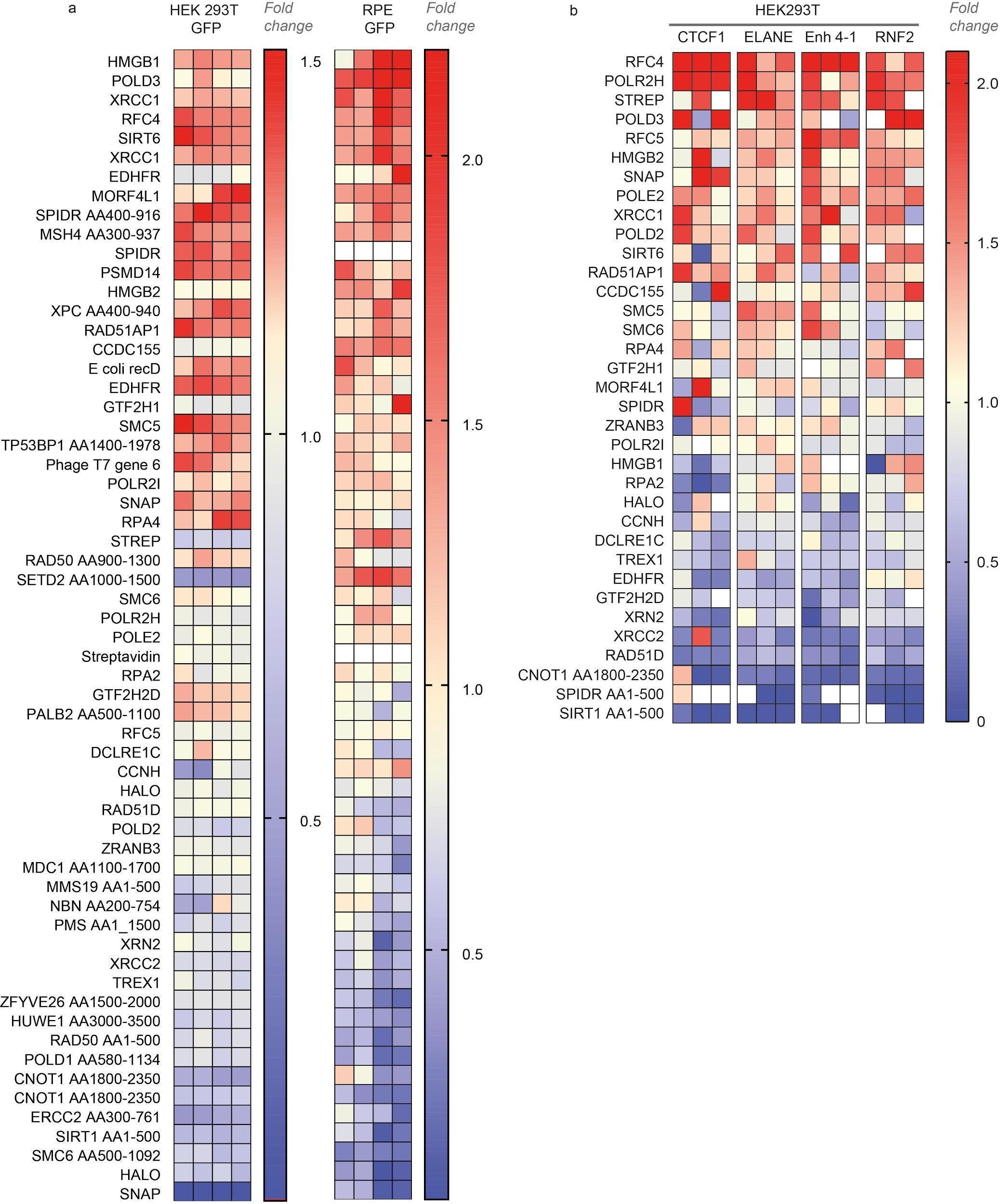

**Figure S2.** **HDR efficiency of the best-performing fusions**.

Each cell in the heatmap illustrates an HDR value that is normalized to the average HDR efficiency of the whole experiment (“fold change”). The sets have an upper limit threshold and values above it are color-coded as a scale maximum (bright-red). For protein fragments, the start and end of the fragment are indicated in relation to the canonical transcript (ie. RAD50 AA900-1300 corresponds to a fragment of RAD50 which starts from amino acid 900 and ends in amino acid 1300). **a**. Normalized GFP conversion rate in HEK293T (n=4 one independent biological experiment) and RPE (n=2, two independent biological experiments) reporter cell lines. **b.** Normalized HDR editing in HEK293T cells that stably express guides targeting endogenous loci (*ELANE, RNF2, Enh4-1* and *CTCF1)* (n=3, one independent biological experiment for each locus). The same dataset is shown in Figure S3 and in Main Figure 1E-F.

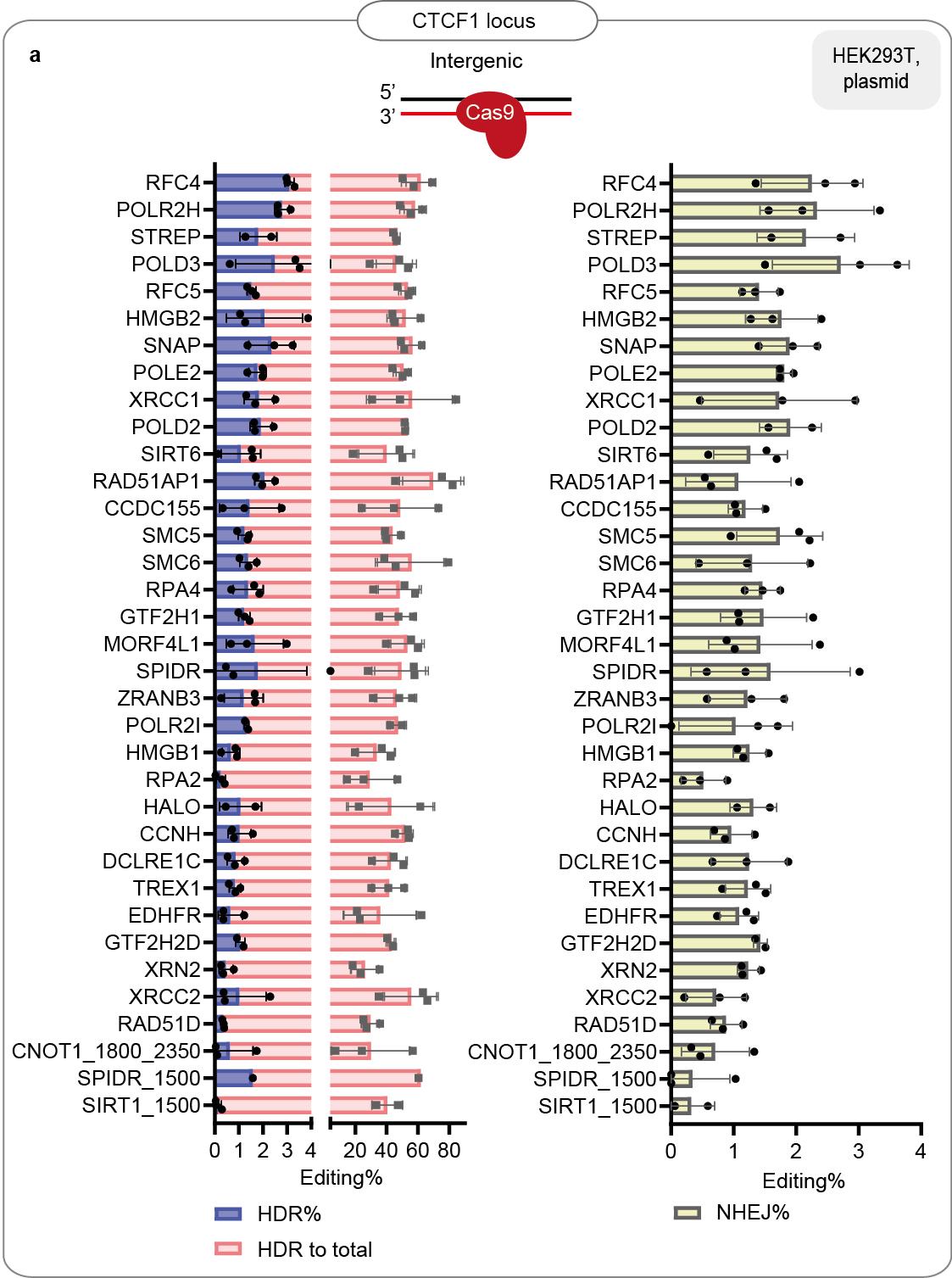

c

c

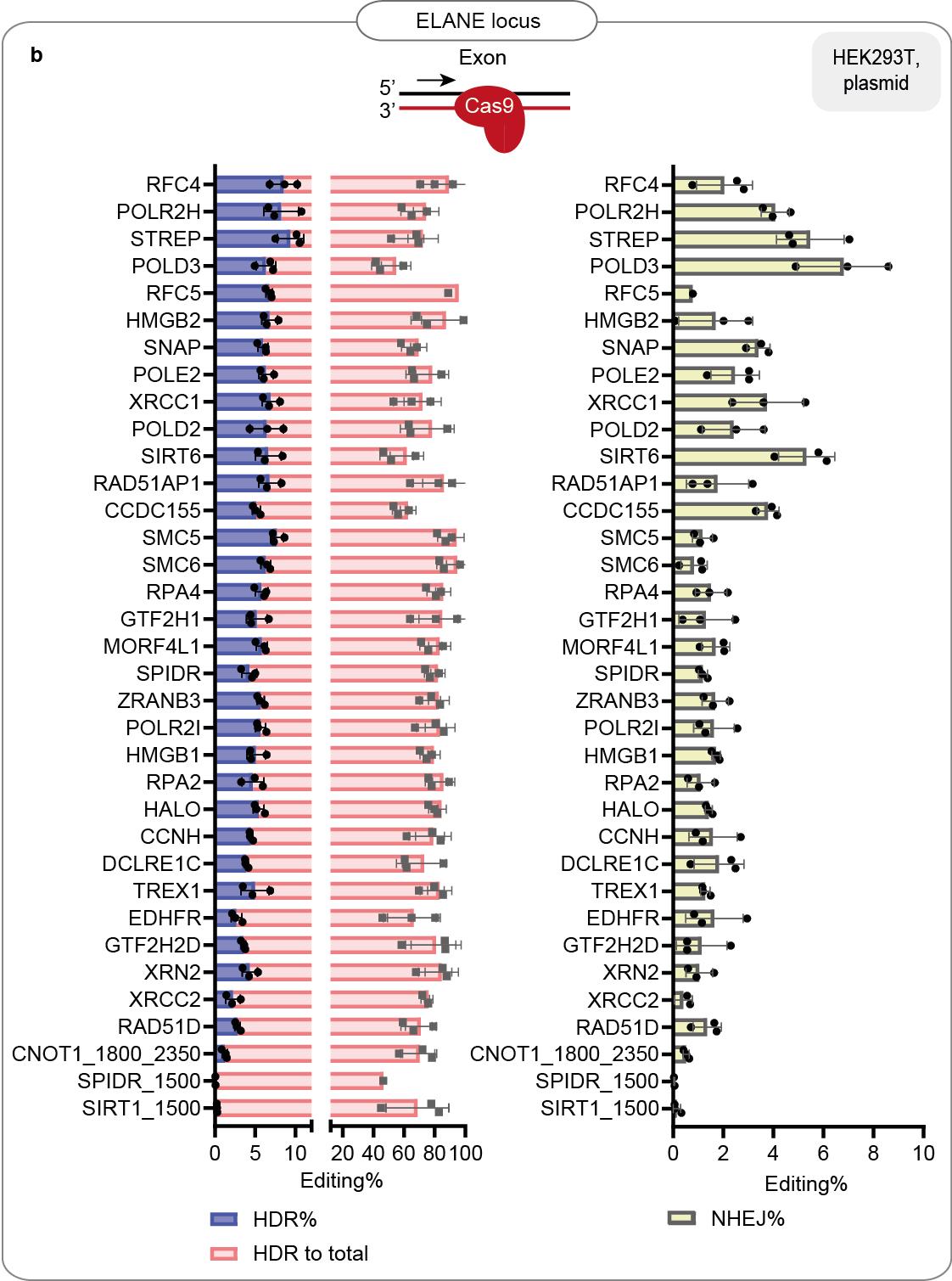

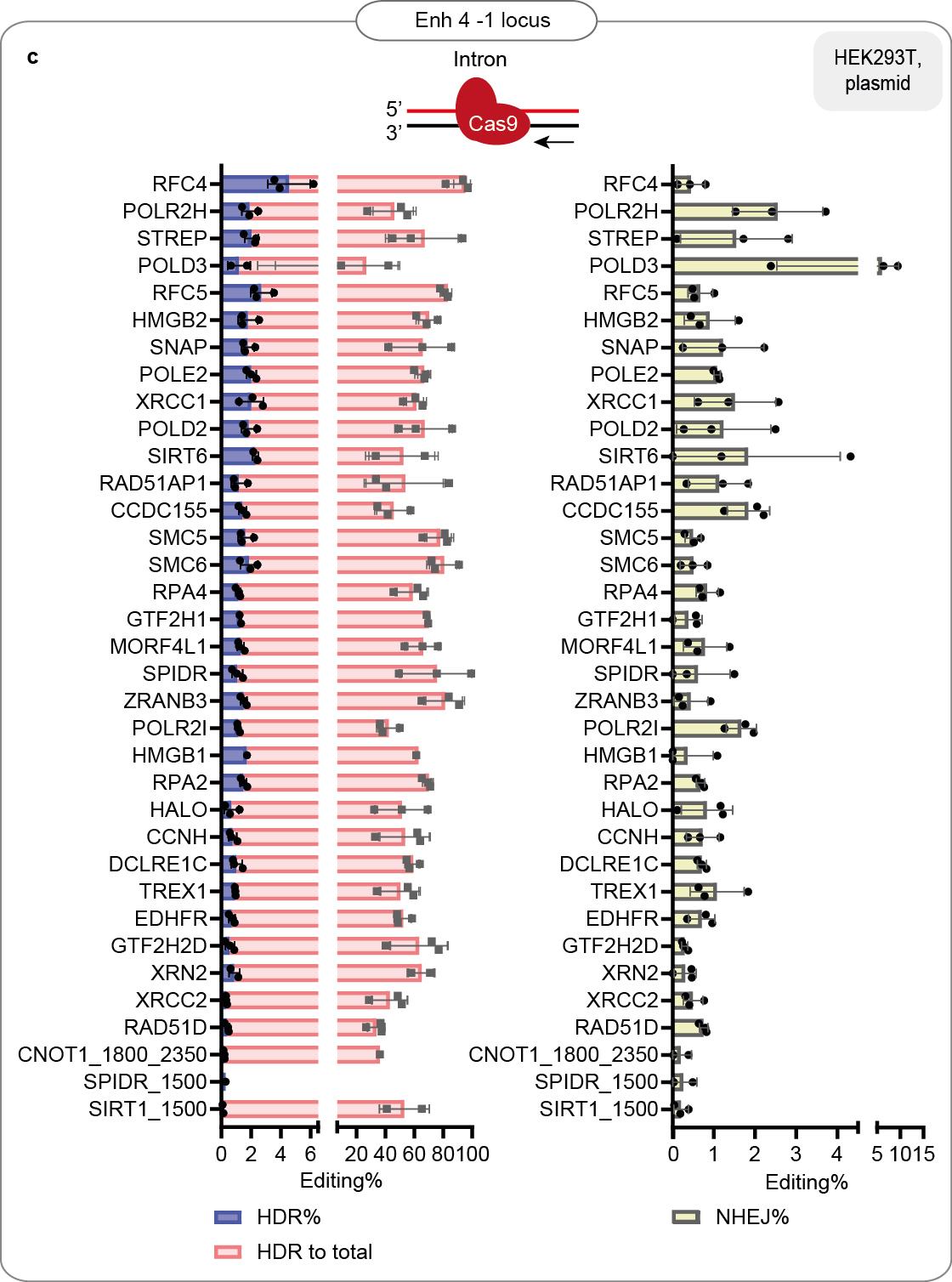

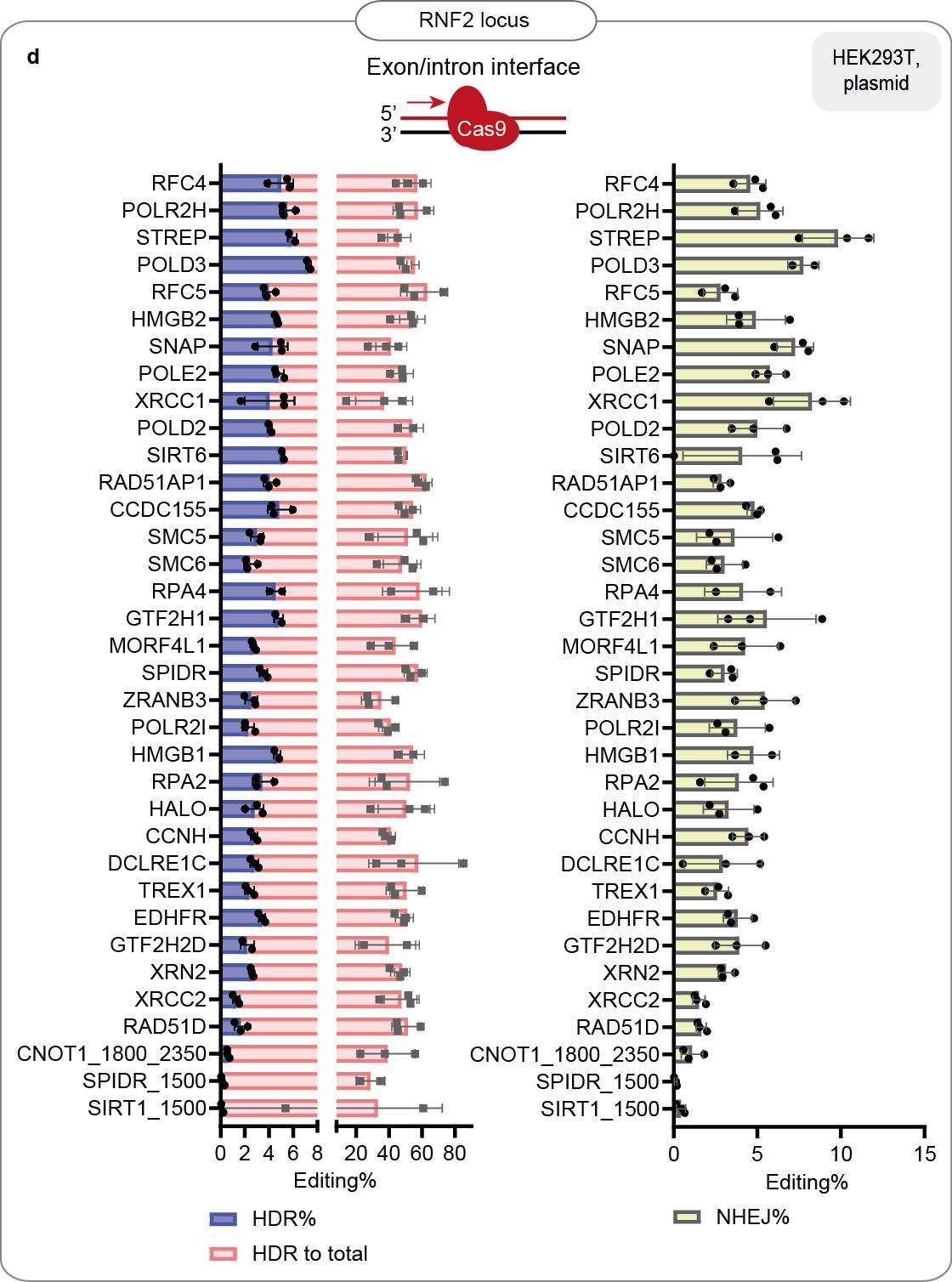

**Figure S3.** **HDR efficiency of the best-performing fusions.** **a-d:** % of HDR, % HDR of total editing and % of NHEJ in HEK293T cells that stably express guides targeting endogenous loci (*CTCF1, ELANE, Enh 4-1* and *RNF2*). Data are the mean values, error bars represent  ±S.D. Top illustrations depict the positioning of the CRISPR-Cas9 complex for each target locus. The red arrow shows the direction of the approaching polymerase. The same dataset is shown in Figure S2B and in Main Figure 1F.

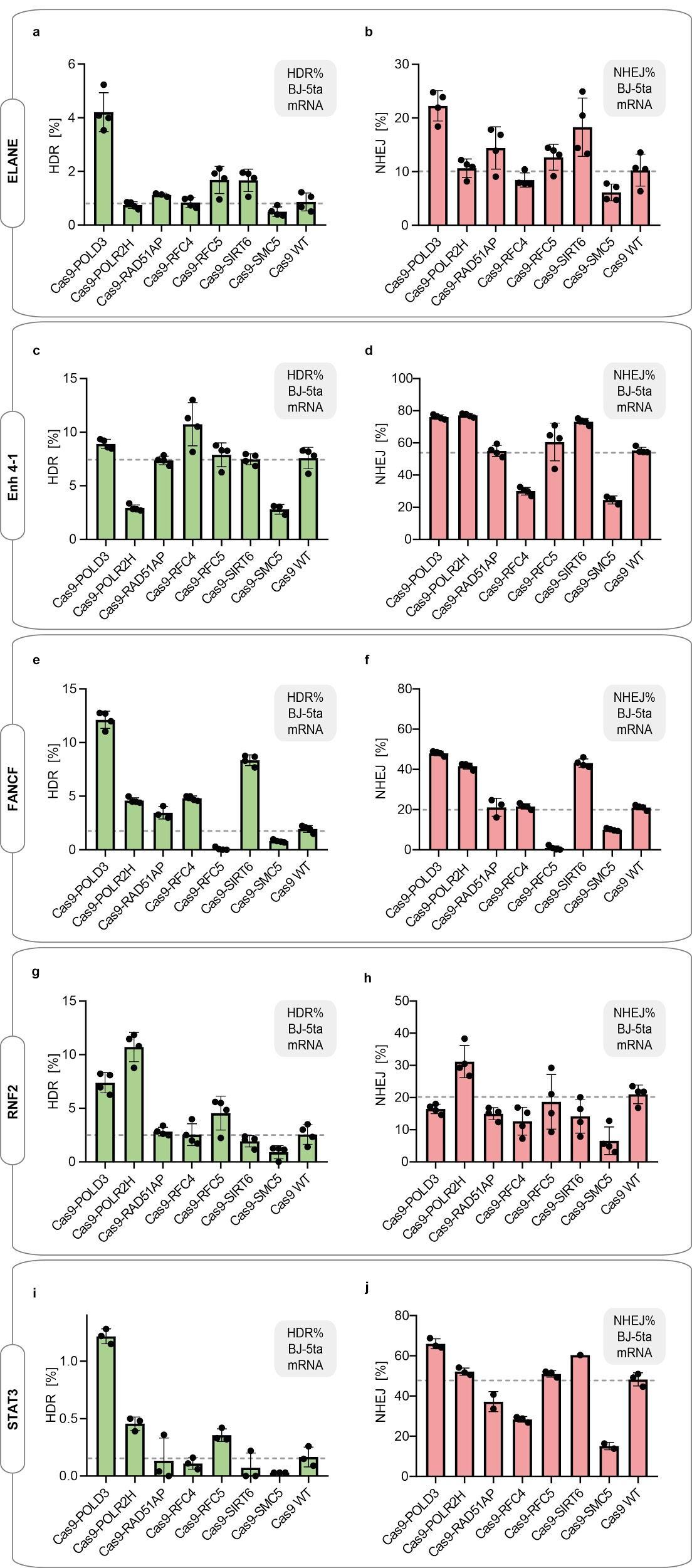

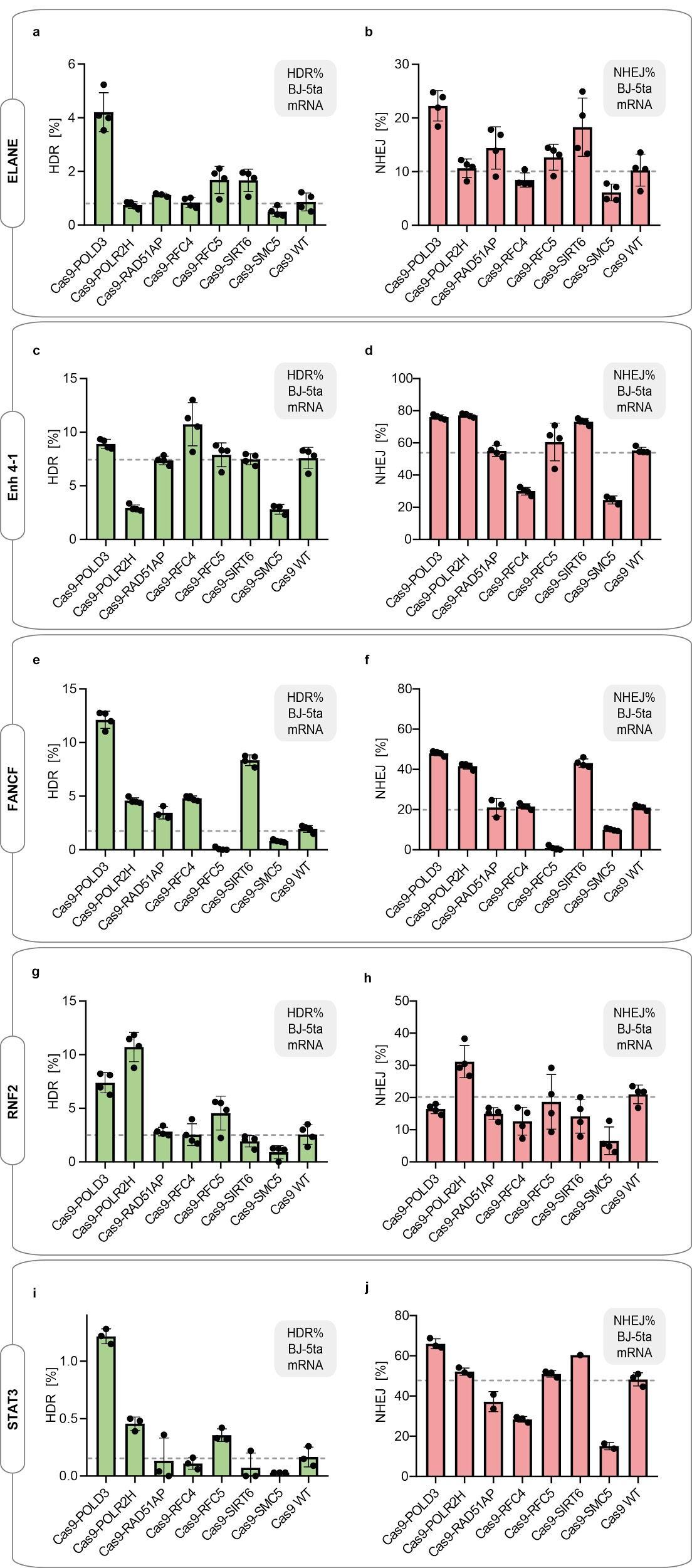
**Figure S4. Cas9 fusion editing efficiency in immortalized fibroblasts (Bj-5ta)** that stably express sgRNA targeting the indicated loci. The cells were electroporated with Cas9 mRNA and repair DNA. HDR and NHEJ editing were measured with ddPCR. n=3, one independent experiment, bar denotes mean value, error bars represent ± S.D. The dataset is also presented in the Main Figure 2B.

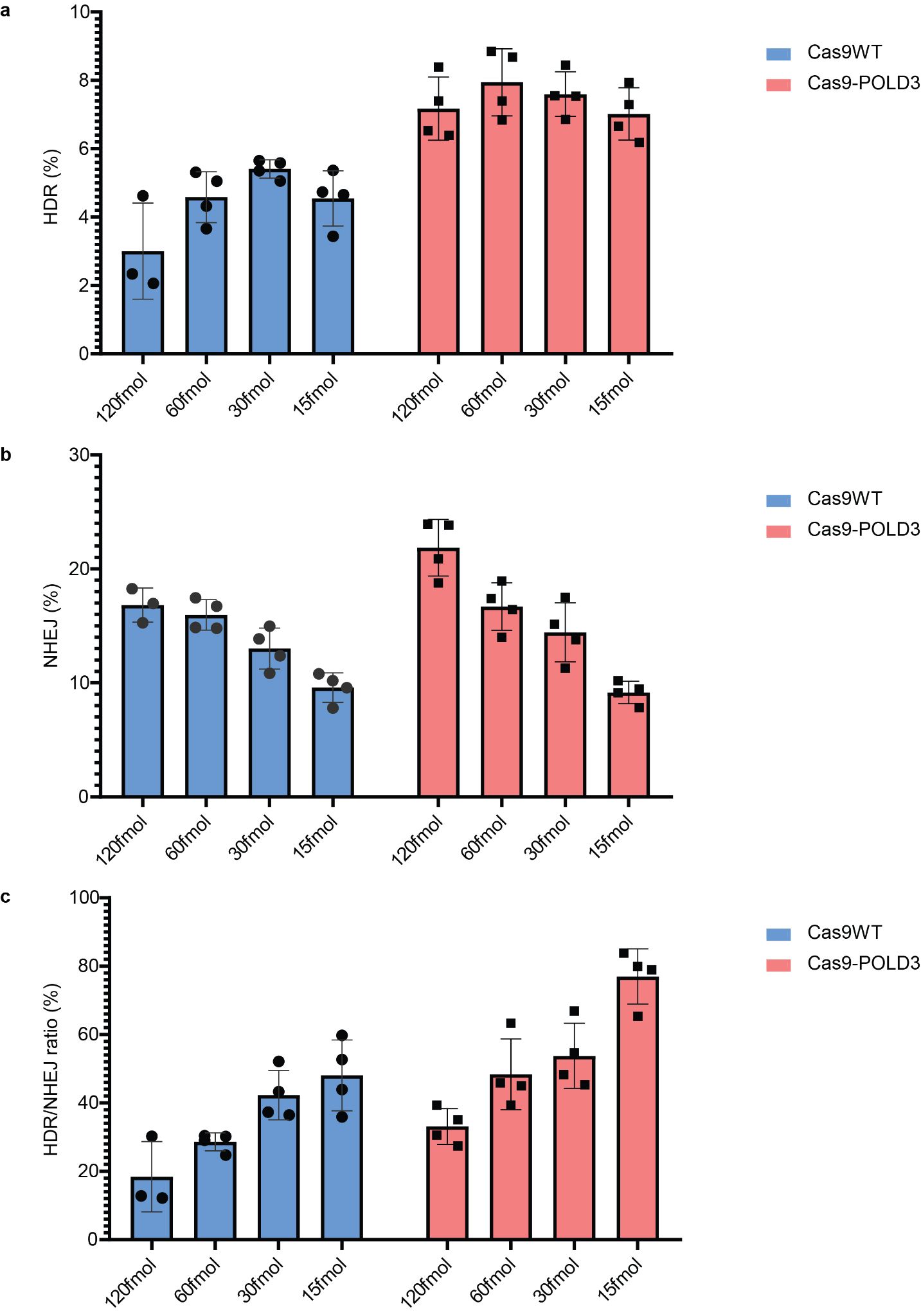

**Figure S5. Cas9-POLD3 editing in decreased concentrations.**

**a-c.** *% of HDR, % NHEJ and % HDR/NHEJ evaluated by ddPCR represents fusion protein performance across different plasmid concentrations in reporter RPE-1 cells, GFP locus. n=4, representative of two independent experiments, bar denotes mean value, error bars represent  ± S.D*

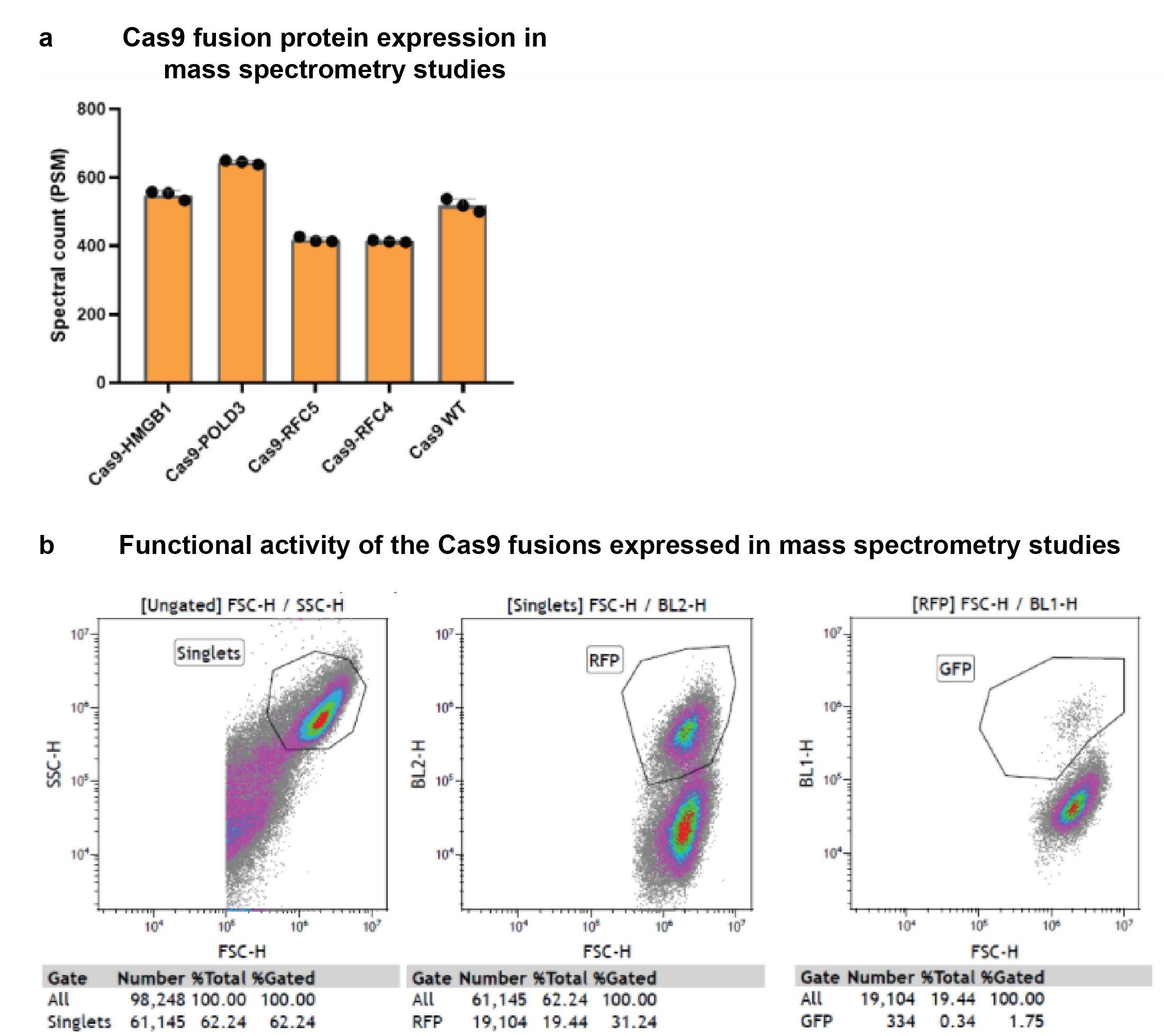
 **Figure S6. Protein expression quality control for mass spectrometry analysis.**

**a.** Peptide-spectrum match (PSM) counts of the Cas9 fusion proteins Flip-in HEK293T cell lines after 30h tetracycline induction. The samples were subsequently used in Affinity Purification Mass Spectrometry (Main Fig.4). **b.** Functional activity of the CRISPR Cas9 fusions used in the mass spectrometry assays. Expression of the fusion construct was induced with tetracycline 24h prior to the transfection with RNP complex targeting GFP locus and corresponding repair template. GFP recovery evaluated by FACS after 5 days.

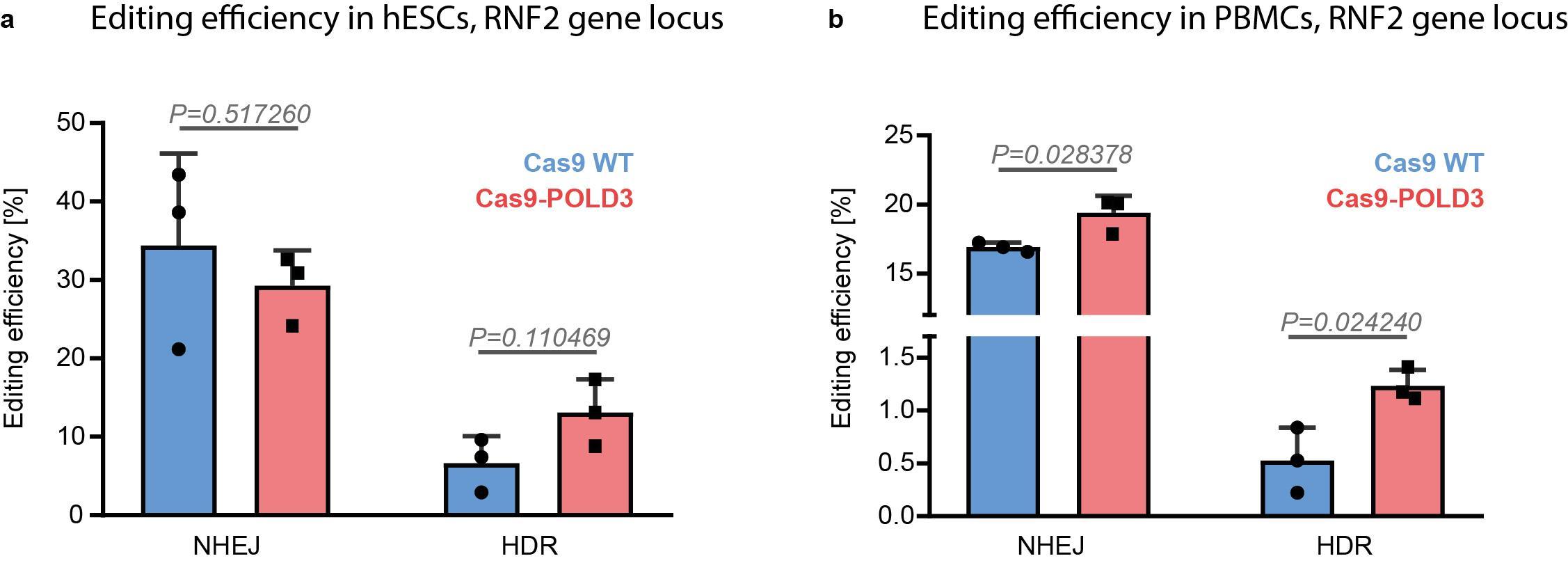

**Figure S7. Cas9-POLD3 editing performance in human embryonic stem cells and peripheral blood mononuclear cells.**

**a.** Human embryonic stem cells (hESCs) and **b.** Peripheral blood mononuclear cells (PBMCs) were electroporated with Cas9 WT and Cas9-POLD3 mRNA, sgRNA targeting the RNF2 locus, and repair DNA. HDR and NHEJ editing were measured with ddPCR. n=3, one independent experiment, bar denotes mean value, error bars represent ± S.D. Statistical significance is calculated with unpaired, two-sided Student’s t-test.

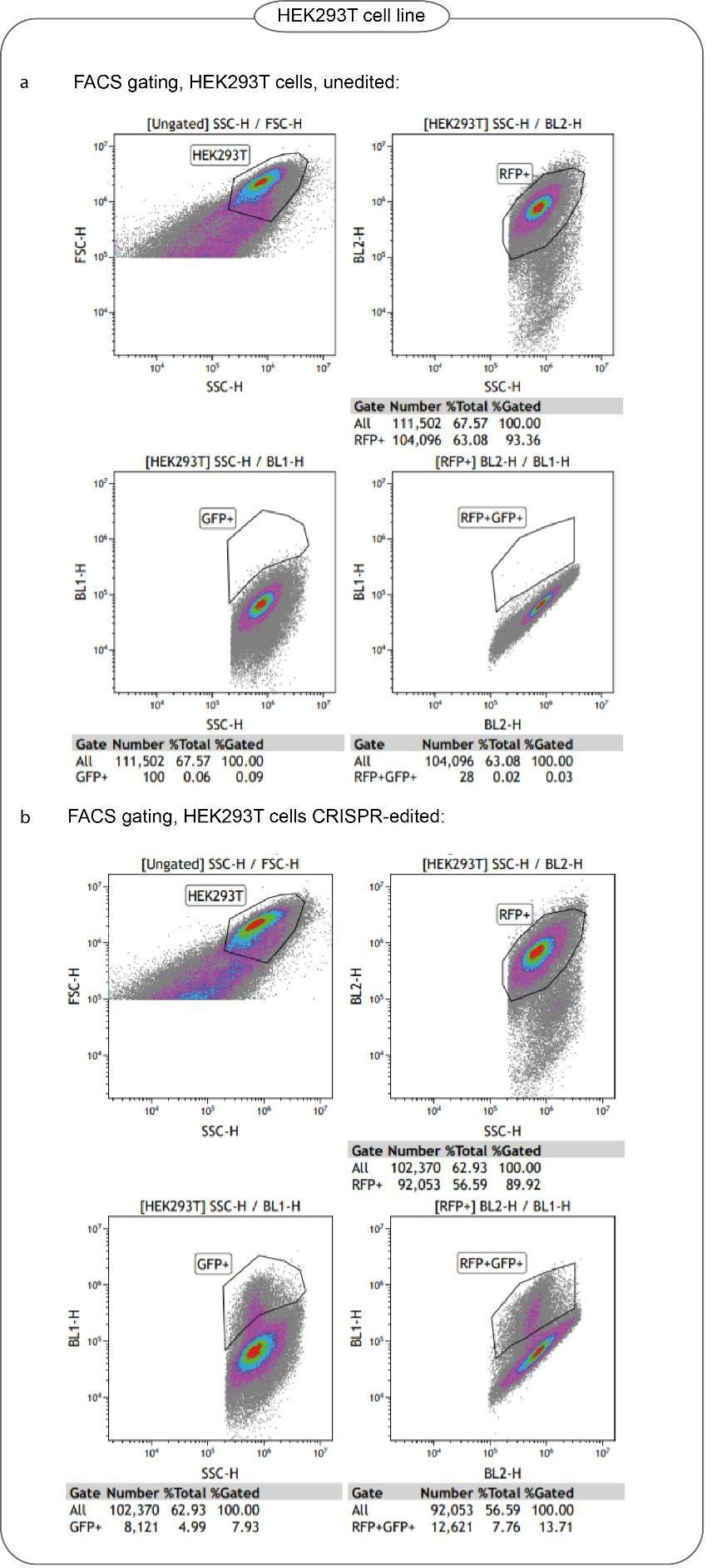

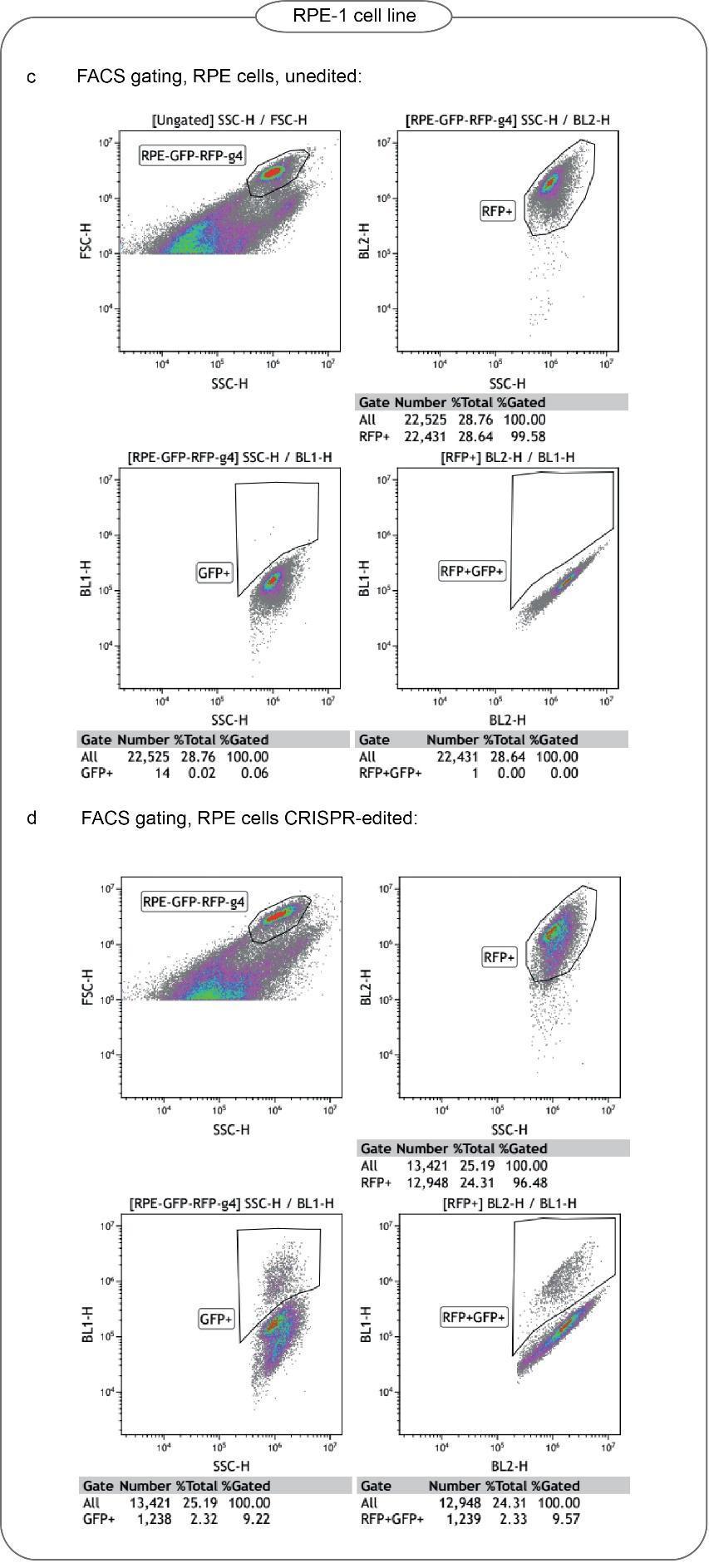

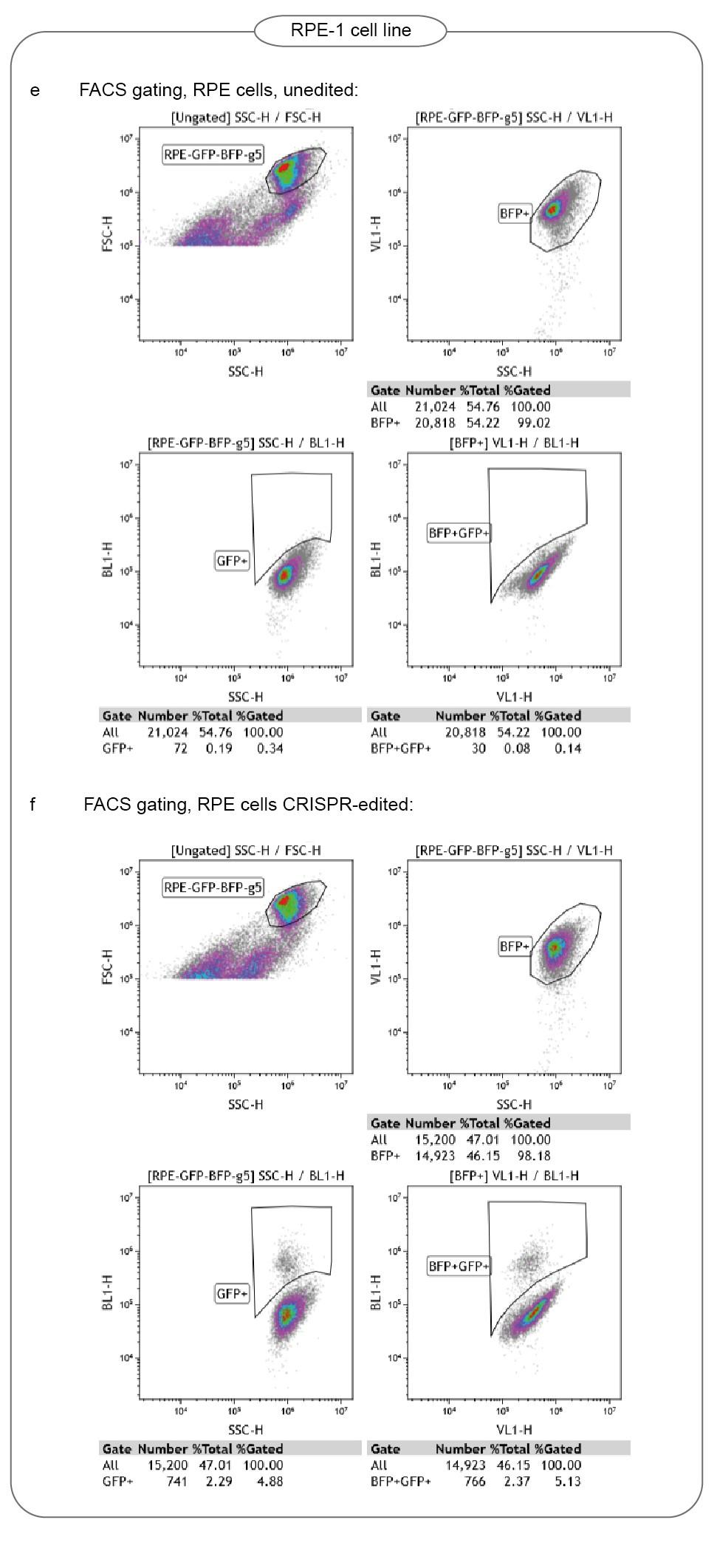

**Figure S8. FACS gating strategy**
**a-b.** HEK293T reporter cell line carrying mGFP-RFP color cassette. **c-d.** RPE-1 reporter cell line carrying mGFP-RFP color cassette. **e-f.** RPE-1 reporter cell line carrying mGFP-BFP color cassette.

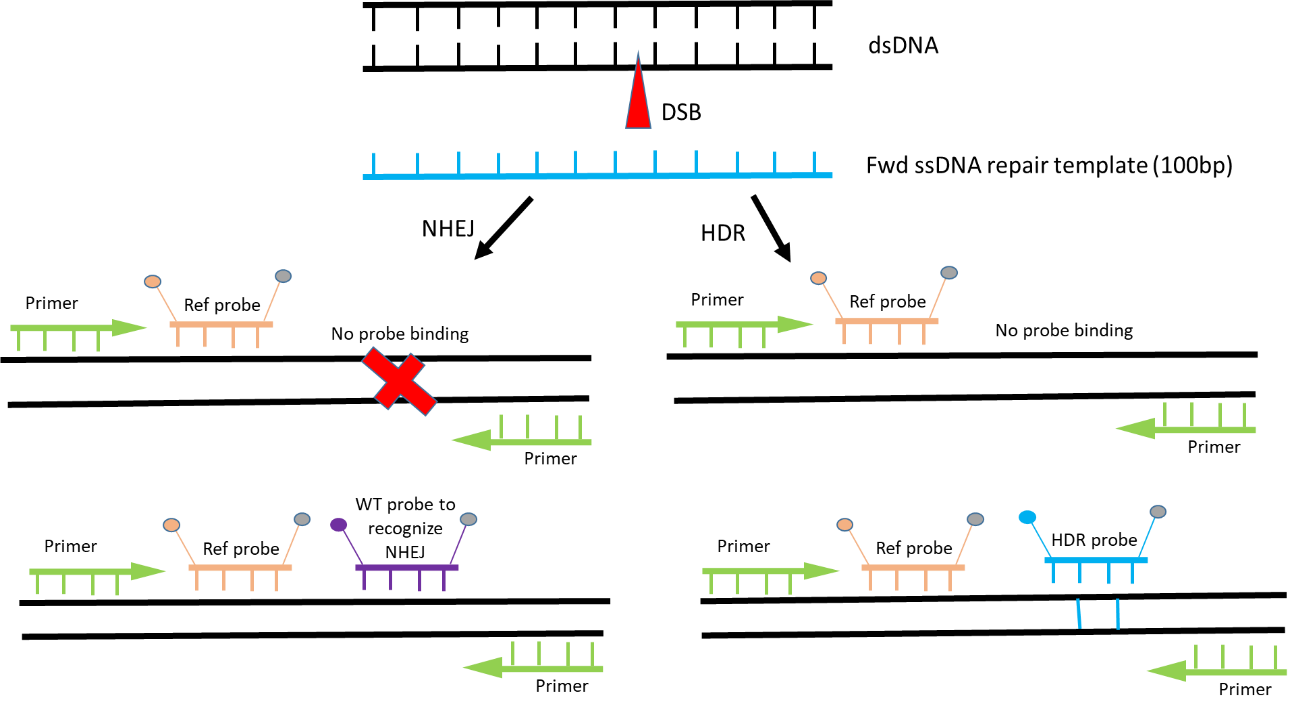

**Figure S9. Quantifying CRISPR editing by droplet digital PCR (ddPCR).**

Primers flank the DNA sequence that surrounds the CRISPR-Cas9 target site. Reference probe binds to the unedited sequence and detects the presence of the amplicon in the droplet. NHEJ probe binds to the WT sequence at the Cas9 target site and drops off if the site mutates because of editing. HDR probe binds to the knock-in DNA sequence from the repair DNA template.

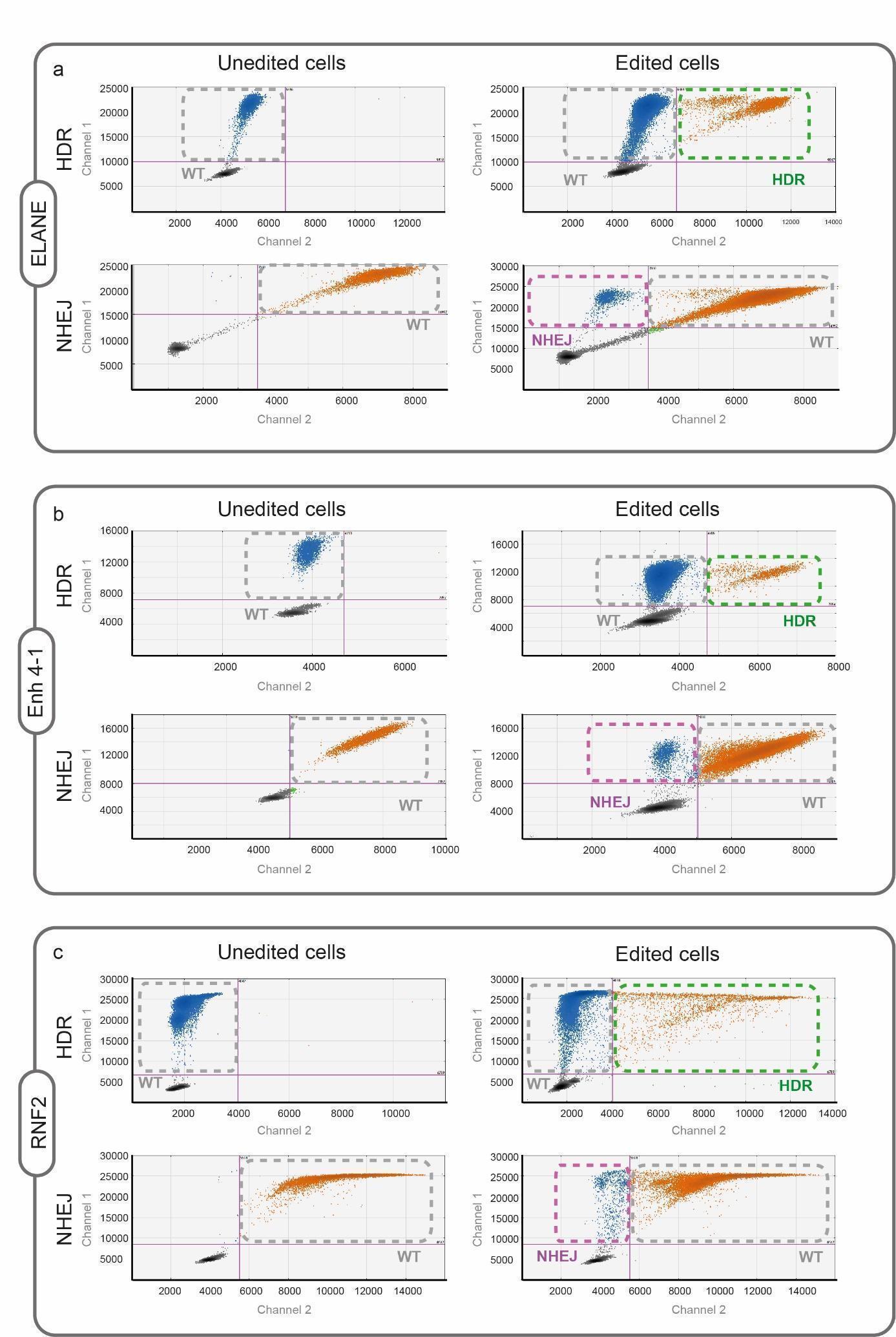

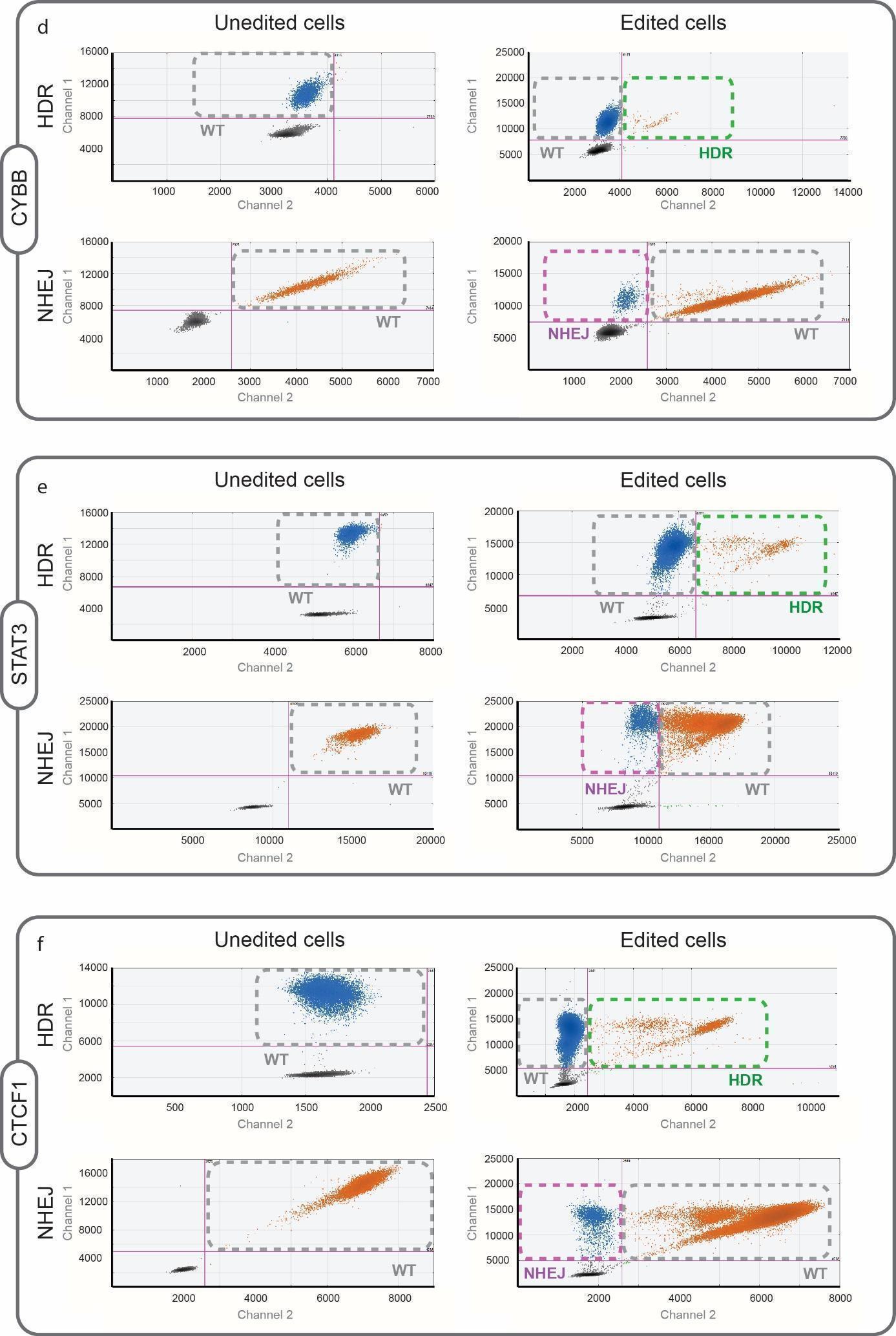

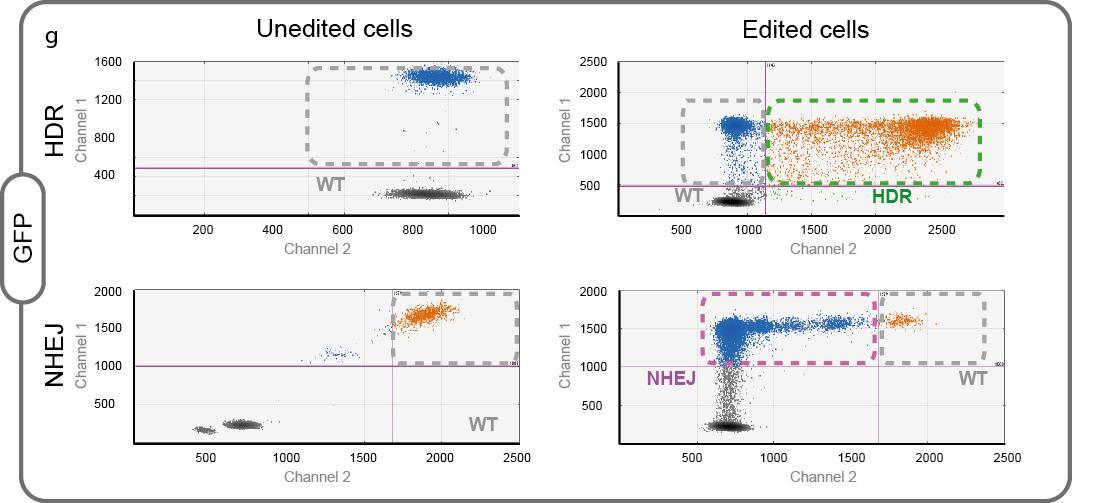
 **Figure S10. Examples of the ddPCR gating for quantifying HDR and NHEJ.**

For each set, the droplet distribution in the negative control is shown on the left and the CRISPR-edited samples on the right.

**Supplementary Table 2:**
Clustering of the hit-fusion’s protein interactions**.** Helicase activity pathways highlighted in orange color.

| **Category** | **Term** | **Count** | **%** | **P value** | **Genes** |
| --- | --- | --- | --- | --- | --- |
| GOTERM_MF_FAT | GO:0003729~mRNA binding | 14 | 12.173913 | 5.84E-10 | P15880, P84103, P16989, P22626, P27797, Q99729, P11940, Q92945, Q14103, O00425, P23396, P62854, P40429, P38159 |
| GOTERM_MF_FAT | GO:0003697~single-stranded DNA binding | 8 | 6.95652174 | 1.01E-05 | P33993, P33991, P10809, P16989, P22626, P61978, Q14566, P15531 |
| GOTERM_MF_FAT | GO:0003678~DNA helicase activity | 6 | 5.2173913 | 5.62E-05 | P33993, P33991, P49736, Q9Y230, Q9Y265, Q14566 |
| GOTERM_MF_FAT | GO:0004003~ATP-dependent DNA helicase activity | 5 | 4.34782609 | 1.70E-04 | P33993, P33991, Q9Y230, Q9Y265, Q14566 |
| GOTERM_MF_FAT | GO:0003677~DNA binding | 34 | 29.5652174 | 6.40E-04 | Q16531, Q9H0D6, P16989, P27797, Q9Y230, P23246, P09874, Q8IUE6, P78527, P0C0S5, P22392, P06899, Q92945, Q9UQ80, P49736, Q00839, P61978, P23396, Q14566, P84090, Q01105, P22626, Q8ND56, P06748, Q99729, Q16777, P15531, P33993, P33991, P16403, P10809, P78347, Q14103, P38159 |
| GOTERM_MF_FAT | GO:0008026~ATP-dependent helicase activity | 6 | 5.2173913 | 0.00106725 | P33993, P33991, Q9Y230, Q9Y265, Q92841, Q14566 |
| GOTERM_MF_FAT | GO:0004386~helicase activity | 7 | 6.08695652 | 0.00114469 | P33993, P33991, P49736, Q9Y230, Q9Y265, Q92841, Q14566 |
| GOTERM_MF_FAT | GO:0008094~DNA-dependent ATPase activity | 5 | 4.34782609 | 0.00300335 | P33993, P33991, Q9Y230, Q9Y265, Q14566 |
| GOTERM_MF_FAT | GO:0042826~histone deacetylase binding | 5 | 4.34782609 | 0.00681963 | P62258, Q14103, P31946, P09874, P23246 |
| GOTERM_MF_FAT | GO:0043139~5'-3' DNA helicase activity | 2 | 1.73913043 | 0.05743824 | Q9Y230, Q9Y265 |
| GOTERM_MF_FAT | GO:0003688~DNA replication origin binding | 2 | 1.73913043 | 0.07812389 | P10809, P49736 |
| GOTERM_MF_FAT | GO:1990837~sequence-specific double-stranded DNA binding | 10 | 8.69565217 | 0.08674837 | P0C0S5, Q9H0D6, P10809, P16989, P49736, Q9Y230, Q14103, P61978, P23246, P15531 |

**Supplementary Table 3:**

CRISPR gRNA sequences of guides that target endogenous loci:

| **Targeted gene** | **Abbreviation** | **Location (Hg19)** | **sgRNA sequence** | **Guide binding strand** | **Gene orientation** |
| --- | --- | --- | --- | --- | --- |
| CCAT1 colon cancer associated transcript 1 | Enh4-1 | chr8:128226403-128226422 | GTAGAATGTCAACTTCATGA | - | REV |
| Transcriptional repressor CTCF | CTCF1 | chr8:128746377-128746396 | TACTTTCGCAAACCTGAACG | + | N/A |
| Ring finger protein 2 | RNF2 | chr1:185056770-185056789 | GTCATCTTAGTCATTACCTG | - | FWD |
| Signal transducer and activator of transcription 3 | STAT3 | chr17:40481574-40481594 | CTCTGCAGAATTCAAACACT | - | REV |
| Elastase, neutrophil expressed | ELANE | chr19:853291-853310 | GAGCCCATAACCTCTCGCGG | + | FWD |
| FANCF FA complementation group F | FANCFA | chr11:2264733522647354 | GGAATCCCTTCTGCAGCACC | - | REV |

**Supplementary Table 4:**
Sequences of ssDNA repair oligos, ddPCR primers and probes:

| **Target gene** | **DNA repair sequence** | **ddPCR primer fwd** | **ddPCR primer rev** | **ddPCR probe NHEJ** | **ddPCR probe HDR** | **ddPCR probe reference** |
| --- | --- | --- | --- | --- | --- | --- |
| **Enh4-1** | ATGTGCTAGACAATATATTAGGTGTGGGATACAGTGAAATCAAATCCTTGagagtcaGAAGTTGACATTCTACTGACAGAAGACATTCAAGCAATACATG | TGTTGAGTGTGCCTGGAT | ACCCTTGCCTCTTTCCCT | CTTGCCCTCATGAAGTTGACATTC | CCTTGagagtcaGAAGTTGACATTCTA | TGGAGGACCGATCATGTGCTAGACA |
| **CTCF-1** | TTTGCTGCAAAGCGTCTTTCCCTCCGCCCCCTCTCTGGGCAGCACttatcaaCAGGTTTGCGAAAGTAAAGTAAGTGTGCCCTCTACTGGCAGCAGAGAT | AGGTGGCTGGAAACTTGT | GGAGCAACCAATCGCTATG | CAAACCTGAACGCGGGTGC | CGCAAACCTGTTGATAAGTGCTGC | CCTCGGACGCTCCTGCTCCT |
| **RNF2** | TAAGCAAAACATGGGAACTCAGTTTATATGAGTTACAACGAACACCTagttcggctcCTAAGATGACTGCCAAGGGGCATATGAGACGTGTAAACTGGGA | CTTCTTTATTTCCAGCAATGTCTC | GCCAACATACAGAAGTCAGG | ACCTCAGGTAATGACTAAGATGACTGC | CTAGTTCGGCTCCTAAGATGACTGC | CTGTGCAGACAAACGGAACTCAA |
| **STAT3** | CCGGCAGCCAGAGGCCCTTTGTGAAGGGGAGCTCCTCCCACATAtaAAcaGTTTGAATTCTGCAatagttCTGCCGTTGTTGGATTCTTCCATGTTCATC | GCTCCCTCAGGGTCTGTA | CCTGTGATTCAGATCCCG | CAAGTGTTTGAATTCTGCAGAGAGGC | TGAATTCTGCAatagttCTGCCGTT | AAAGGCAGGTGTCCTGTGA |
| **ELANE** | GTTTTCGAAGATGCGCTGCACGGCGAACACCTGCCGGGTGGGCTCttGatGtGAGAGGTTATGGGCTCCCAGGACCACCCGCACCGCGCGGACGTTTCTG | GGATGGGGACGACAAGG | GCACCTGGAGAATCACGA | CTCGCGGCGGGAGC | CATAACCTCTCaCatCaaGAGCCC | ACCCCGTAAACTTGCTCAACGA |
| **FANCF** | GGTGCTGACGTAGGTAGTGCTTGAGACCGCCAGAAGCTCGGAAAAGCGATtgtatcGCTGCAGAAGGGATTCCATGAGGTGCGCGAAGGCCCTACTTCCG | ATGGATGTGGCGCAGGTA | CGCGGATGTTCCAATCAG | CGATCCAGGTGCTGCAGAA | AGCGATtgtatcGCTGCAGAA | TCACCTTGGAGACGGCGAC |
| **GFP** | GAACGGCCACAAGTTCAGCGTGTCCGGCGAGGGCGAAGGCGACGCCACATATGGCAAGCTGACCCTGAAGTTCATCTGCACCACCGGCAAGCTGCCCGTG | ATCCTGGTGGAACTGGATG | GAAGTCGTGCTGCTTCATAT | GCCGCTGCCAAGCTGAC | CGCCACATATGGCAAGCTGAC | GCGACGTGAACGGCCAC |

**Letters written in non-capital font in the “DNA repair sequence” column represent nucleotides that are different from the original WT gene sequence.*

**Appendix I**

Cas9-GW sequence:

Nuclear localization signal

Cas9 sequence

GS linker

ATGAAAAGGCCGGCGGCCACGAAAAAGGCCGGCCAGGCAAAAAAGAAAAAGGACAAGAAGTACAGCATCGGCCTGGACATCGGCACCAACTCTGTGGGCTGGGCCGTGATCACCGACGAGTACAAGGTGCCCAGCAAGAAATTCAAGGTGCTGGGCAACACCGACCGGCACAGCATCAAGAAGAACCTGATCGGAGCCCTGCTGTTCGACAGCGGCGAAACAGCCGAGGCCACCCGGCTGAAGAGAACCGCCAGAAGAAGATACACCAGACGGAAGAACCGGATCTGCTATCTGCAAGAGATCTTCAGCAACGAGATGGCCAAGGTGGACGACAGCTTCTTCCACAGACTGGAAGAGTCCTTCCTGGTGGAAGAGGATAAGAAGCACGAGCGGCACCCCATCTTCGGCAACATCGTGGACGAGGTGGCCTACCACGAGAAGTACCCCACCATCTACCACCTGAGAAAGAAACTGGTGGACAGCACCGACAAGGCCGACCTGCGGCTGATCTATCTGGCCCTGGCCCACATGATCAAGTTCCGGGGCCACTTCCTGATCGAGGGCGACCTGAACCCCGACAACAGCGACGTGGACAAGCTGTTCATCCAGCTGGTGCAGACCTACAACCAGCTGTTCGAGGAAAACCCCATCAACGCCAGCGGCGTGGACGCCAAGGCCATCCTGTCTGCCAGACTGAGCAAGAGCAGACGGCTGGAAAATCTGATCGCCCAGCTGCCCGGCGAGAAGAAGAATGGCCTGTTCGGCAACCTGATTGCCCTGAGCCTGGGCCTGACCCCCAACTTCAAGAGCAACTTCGACCTGGCCGAGGATGCCAAACTGCAGCTGAGCAAGGACACCTACGACGACGACCTGGACAACCTGCTGGCCCAGATCGGCGACCAGTACGCCGACCTGTTTCTGGCCGCCAAGAACCTGTCCGACGCCATCCTGCTGAGCGACATCCTGAGAGTGAACACCGAGATCACCAAGGCCCCCCTGAGCGCCTCTATGATCAAGAGATACGACGAGCACCACCAGGACCTGACCCTGCTGAAAGCTCTCGTGCGGCAGCAGCTGCCTGAGAAGTACAAAGAGATTTTCTTCGACCAGAGCAAGAACGGCTACGCCGGCTACATTGACGGCGGAGCCAGCCAGGAAGAGTTCTACAAGTTCATCAAGCCCATCCTGGAAAAGATGGACGGCACCGAGGAACTGCTCGTGAAGCTGAACAGAGAGGACCTGCTGCGGAAGCAGCGGACCTTCGACAACGGCAGCATCCCCCACCAGATCCACCTGGGAGAGCTGCACGCCATTCTGCGGCGGCAGGAAGATTTTTACCCATTCCTGAAGGACAACCGGGAAAAGATCGAGAAGATCCTGACCTTCCGCATCCCCTACTACGTGGGCCCTCTGGCCAGGGGAAACAGCAGATTCGCCTGGATGACCAGAAAGAGCGAGGAAACCATCACCCCCTGGAACTTCGAGGAAGTGGTGGACAAGGGCGCTTCCGCCCAGAGCTTCATCGAGCGGATGACCAACTTCGATAAGAACCTGCCCAACGAGAAGGTGCTGCCCAAGCACAGCCTGCTGTACGAGTACTTCACCGTGTATAACGAGCTGACCAAAGTGAAATACGTGACCGAGGGAATGAGAAAGCCCGCCTTCCTGAGCGGCGAGCAGAAAAAGGCCATCGTGGACCTGCTGTTCAAGACCAACCGGAAAGTGACCGTGAAGCAGCTGAAAGAGGACTACTTCAAGAAAATCGAGTGCTTCGACTCCGTGGAAATCTCCGGCGTGGAAGATCGGTTCAACGCCTCCCTGGGCACATACCACGATCTGCTGAAAATTATCAAGGACAAGGACTTCCTGGACAATGAGGAAAACGAGGACATTCTGGAAGATATCGTGCTGACCCTGACACTGTTTGAGGACAGAGAGATGATCGAGGAACGGCTGAAAACCTATGCCCACCTGTTCGACGACAAAGTGATGAAGCAGCTGAAGCGGCGGAGATACACCGGCTGGGGCAGGCTGAGCCGGAAGCTGATCAACGGCATCCGGGACAAGCAGTCCGGCAAGACAATCCTGGATTTCCTGAAGTCCGACGGCTTCGCCAACAGAAACTTCATGCAGCTGATCCACGACGACAGCCTGACCTTTAAAGAGGACATCCAGAAAGCCCAGGTGTCCGGCCAGGGCGATAGCCTGCACGAGCACATTGCCAATCTGGCCGGCAGCCCCGCCATTAAGAAGGGCATCCTGCAGACAGTGAAGGTGGTGGACGAGCTgGTGAAAGTGATGGGCCGGCACAAGCCCGAGAACATCGTGATCGAAATGGCCAGAGAGAACCAGACCACCCAGAAGGGACAGAAGAACAGCCGCGAGAGAATGAAGCGGATCGAAGAGGGCATCAAAGAGCTGGGCAGCCAGATCCTGAAAGAACACCCCGTGGAAAACACCCAGCTGCAGAACGAGAAGCTGTACCTGTACTACCTGCAGAATGGGCGGGATATGTACGTGGACCAGGAACTGGACATCAACCGGCTGTCCGACTACGATGTGGACCACATCGTGCCTCAGAGCTTTCTGAAGGACGACTCCATCGACAACAAGGTGCTGACCAGAAGCGACAAgAACCGGGGCAAGAGCGACAACGTGCCCTCCGAAGAGGTCGTGAAGAAGATGAAGAACTACTGGCGGCAGCTGCTGAACGCCAAGCTGATTACCCAGAGAAAGTTCGACAATCTGACCAAGGCCGAGAGAGGCGGCCTGAGCGAACTGGATAAGGCCGGCTTCATCAAGAGACAGCTGGTGGAAACCCGGCAGATCACAAAGCACGTGGCACAGATCCTGGACTCCCGGATGAACACTAAGTACGACGAGAATGACAAGCTGATCCGGGAAGTGAAAGTGATCACCCTGAAGTCCAAGCTGGTGTCCGATTTCCGGAAGGATTTCCAGTTTTACAAAGTGCGCGAGATCAACAACTACCACCACGCCCACGACGCCTACCTGAACGCCGTCGTGGGAACCGCCCTGATCAAAAAGTACCCTAAGCTGGAAAGCGAGTTCGTGTACGGCGACTACAAGGTGTACGACGTGCGGAAGATGATCGCCAAGAGCGAGCAGGAAATCGGCAAGGCTACCGCCAAGTACTTCTTCTACAGCAACATCATGAACTTTTTCAAGACCGAGATTACCCTGGCCAACGGCGAGATCCGGAAGCGGCCTCTGATCGAGACAAACGGCGAAACCGGGGAGATCGTGTGGGATAAGGGCCGGGATTTTGCCACCGTGCGGAAAGTGCTGAGCATGCCCCAAGTGAATATCGTGAAAAAGACCGAGGTGCAGACAGGCGGCTTCAGCAAAGAGTCTATCCTGCCCAAGAGGAACAGCGATAAGCTGATCGCCAGAAAGAAGGACTGGGACCCTAAGAAGTACGGCGGCTTCGACAGCCCCACCGTGGCCTATTCTGTGCTGGTGGTGGCCAAAGTGGAAAAGGGCAAGTCCAAGAAACTGAAGAGTGTGAAAGAGCTGCTGGGGATCACCATCATGGAAAGAAGCAGCTTCGAGAAGAATCCCATCGACTTTCTGGAAGCCAAGGGCTACAAAGAAGTGAAAAAGGACCTGATCATCAAGCTGCCTAAGTACTCCCTGTTCGAGCTGGAAAACGGCCGGAAGAGAATGCTGGCCTCTGCCGGCGAACTGCAGAAGGGAAACGAACTGGCCCTGCCCTCCAAATATGTGAACTTCCTGTACCTGGCCAGCCACTATGAGAAGCTGAAGGGCTCCCCCGAGGATAATGAGCAGAAACAGCTGTTTGTGGAACAGCACAAGCACTACCTGGACGAGATCATCGAGCAGATCAGCGAGTTCTCCAAGAGAGTGATCCTGGCCGACGCTAATCTGGACAAAGTGCTGTCCGCCTACAACAAGCACCGGGATAAGCCCATCAGAGAGCAGGCCGAGAATATCATCCACCTGTTTACCCTGACCAATCTGGGAGCCCCTGCCGCCTTCAAGTACTTTGACACCACCATCGACCGGAAGAGGTACACCAGCACCAAAGAGGTGCTGGACGCCACCCTGATCCACCAGAGCATCACCGGCCTGTACGAGACACGGATCGACCTGTCTCAGCTGGGAGGCGACagcgctGGAGGAGGTGGAAGCGGAGGAGGAGGAAGCGGAGGAGGAGGTAGCggacctaagaaaaagaggaaggtggGAATTC-GW recombination site – TAA

**Appendix II**

GFP-BFP reporter cassette sequence:

LTR
U6
gRNA for mutant GFP

Kozak
Zeocin
GS linker
PAM
Optimised emGFP
GFP gRNA binding site

2A peptide
BFP

gggtctctctggttagaccagatctgagcctgggagctctctggctaactagggaacccactgcttaagcctcaataaagcttgccttgagtgcttcaagtagtgtgtgcccgtctgttgtgtgactctggtaactagagatccctcagacccttttagtcagtgtggaaaatctctagcagtggcgcccgaacagggacttgaaagcgaaagggaaaccagaggagctctctcgacgcaggactcggcttgctgaagcgcgcacggcaagaggcgaggggcggcgactggtgagtacgccaaaaattttgactagcggaggctagaaggagagagatgggtgcgagagcgtcagtattaagcgggggagaattagatcgcgatgggaaaaaattcggttaaggccagggggaaagaaaaaatataaattaaaacatatagtatgggcaagcagggagctagaacgattcgcagttaatcctggcctgttagaaacatcagaaggctgtagacaaatactgggacagctacaaccatcccttcagacaggatcagaagaacttagatcattatataatacagtagcaaccctctattgtgtgcatcaaaggatagagataaaagacaccaaggaagctttagacaagatagaggaagagcaaaacaaaagtaagaccaccgcacagcaagcggccgctgatcttcagacctggaggaggagatatgagggacaattggagaagtgaattatataaatataaagtagtaaaaattgaaccattaggagtagcacccaccaaggcaaagagaagagtggtgcagagagaaaaaagagcagtgggaataggagctttgttccttgggttcttgggagcagcaggaagcactatgggcgcagcgtcaatgacgctgacggtacaggccagacaattattgtctggtatagtgcagcagcagaacaatttgctgagggctattgaggcgcaacagcatctgttgcaactcacagtctggggcatcaagcagctccaggcaagaatcctggctgtggaaagatacctaaaggatcaacagctcctggggatttggggttgctctggaaaactcatttgcaccactgctgtgccttggaatgctagttggagtaataaatctctggaacagatttggaatcacacgacctggatggagtgggacagagaaattaacaattacacaagcttaatacactccttaattgaagaatcgcaaaaccagcaagaaaagaatgaacaagaattattggaattagataaatgggcaagtttgtggaattggtttaacataacaaattggctgtggtatataaaattattcataatgatagtaggaggcttggtaggtttaagaatagtttttgctgtactttctatagtgaatagagttaggcagggatattcaccattatcgtttcagacccacctcccaaccccgaggggacccgacaggcccgaaggaatagaagaagaaggtggagagagagacagagacagatccattcgattagtgaacggatcggcactgcgtgcgccaattctgcagacaaatggcagtattcatccacaattttaaaagaaaaggggggattggggggtacagtgcaggggaaagaatagtagacataatagcaacagacatacaaactaaagaattacaaaaacaaattacaaaaattcaaaattttcgggtttattacagggacagcagagatccagtttggttaattagctagcgagggcctatttcccatgattccttcatatttgcatatacgatacaaggctgttagagagataattggaattaatttgactgtaaacacaaagatattagtacaaaatacgtgacgtagaaagtaataatttcttgggtagtttgcagttttaaaattatgttttaaaatggactatcatatgcttaccgtaacttgaaagtatttcgatttcttggctttatatatcttGTGGAAAGGACGAAACACCgGAGACGggatacCGTCTCtgttttagagctaggccAACATGAGGATCACCCATGTCTGCAGggcctagcaagttaaaataaggctagtccgttatcaacttggccAACATGAGGATCACCCATGTCTGCAGggccaagtggcaccgagtcggtgcTTTTTTTggatcctgcaaagatggataaagttttaaacagagaggaatctttgcagctaatggaccttctaggtcttgaaaggagtgggaattggctccggtgcccgtcagtgggcagagcgcacatcgcccacagtccccgagaagttggggggaggggtcggcaattgatccggtgcctagagaaggtggcgcggggtaaactgggaaagtgatgtcgtgtactggctccgcctttttcccgagggtgggggagaaccgtatataagtgcagtagtcgccgtgaacgttctttttcgcaacgggtttgccgccagaacacaggtaagtgccgtgtgtggttcccgcgggcctggcctctttacgggttatggcccttgcgtgccttgaattacttccactggctgcagtacgtgattcttgatcccgagcttcgggttggaagtgggtgggagagttcgaggccttgcgcttaaggagccccttcgcctcgtgcttgagttgaggcctggcctgggcgctggggccgccgcgtgcgaatctggtggcaccttcgcgcctgtctcgctgctttcgataagtctctagccatttaaaatttttgatgacctgctgcgacgctttttttctggcaagatagtcttgtaaatgcgggccaagatctgcacactggtatttcggtttttggggccgcgggcggcgacggggcccgtgcgtcccagcgcacatgttcggcgaggcggggcctgcgagcgcggccaccgagaatcggacgggggtagtctcaagctggccggcctgctctggtgcctggcctcgcgccgccgtgtatcgccccgccctgggcggcaaggctggcccggtcggcaccagttgcgtgagcggaaagatggccgcttcccggccctgctgcagggagctcaaaatggaggacgcggcgctcgggagagcgggcgggtgagtcacccacacaaaggaaaagggcctttccgtcctcagccgtcgcttcatgtgactccacggagtaccgggcgccgtccaggcacctcgattagttctcgagcttttggagtacgtcgtctttaggttggggggaggggttttatgcgatggagtttccccacactgagtgggtggagactgaagttaggccagcttggcacttgatgtaattctccttggaatttgccctttttgagtttggatcttggttcattctcaagcctcagacagtggttcaaagtttttttcttccatttcaggtgtcgtgatgtacatgccaccATGGCCAAGTTGACCAGTGCCGTTCCGGTGCTCACCGCGCGCGACGTCGCCGGAGCGGTCGAGTTCTGGACCGACCGGCTCGGGTTCTCCCGGGACTTCGTGGAGGACGACTTCGCCGGTGTGGTCCGGGACGACGTGACCCTGTTCATCAGCGCGGTCCAGGACCAGGTGGTGCCGGACAACACCCTGGCCTGGGTGTGGGTGCGCGGCCTGGACGAGCTGTACGCCGAGTGGTCGGAGGTCGTGTCCACGAACTTCCGGGACGCCTCCGGGCCGGCCATGACCGAGATCGGCGAGCAGCCGTGGGGGCGGGAGTTCGCCCTGCGCGACCCGGCCGGCAACTGCGTGCACTTCGTGGCCGAGGAGCAGGACGGAGGAGGTGGAAGCGGAGGAGGAGGAAGCGTGAGCAAGGGCGAGGAACTGTTCACCGGCGTGGTGCCCATCCTGGTGGAACTGGATGGCGACGTGAACGGCCACAAGTTCAGCGTGTCCGGCGAGGGCGAAGGCGACGCCGCCGCTGCCAAGCTGACCCTGAAGTTCATCTGCACCACCGGCAAGCTGCCCGTGCCTTGGCCTACCCTCGTGACCACCTTTACCTACGGCGTGCAGTGCTTCGCCAGATACCCCGACCATATGAAGCAGCACGACTTCTTCAAGAGCGCCATGCCCGAGGGCTACGTGCAGGAACGGACCATCTTCTTTAAGGACGACGGCAACTACAAGACCAGGGCCGAAGTGAAGTTCGAGGGCGACACCCTCGTGAACCGGATCGAGCTGAAGGGCATCGACTTCAAAGAGGACGGCAACATCCTGGGCCACAAGCTGGAGTACAACTACAACAGCCACAAGGTcTACATCACCGCCGACAAGCAGAAAAACGGCATCAAAGTGAACTTCAAGACCCGGCACAACATCGAGGACGGCTCCGTGCAGCTGGCCGACCACTACCAGCAGAACACCCCCATCGGAGATGGCCCCGTGCTGCTGCCCGACAACCACTACCTGAGCACCCAGAGCAAGCTGAGCAAGGACCCCAACGAGAAGCGGGACCACATGGTGCTGCTGGAATTTGTGACCGCCGCTGGCATCACCCTGGGCATGGACGAGCTcTACAAGGGATCCGGCGCAACAAACTTCTCTCTGCTGAAACAAGCCGGAGATGTCGAAGAGAATCCTGGACCGAGCGAGCTGATTAAGGAGAACATGCACATGAAGCTcTACATGGAGGGCACCGTGGACAACCATCACTTCAAGTGCACATCCGAGGGCGAAGGCAAGCCCTACGAGGGCACCCAGACCATGAGAATCAAGGTGGTCGAGGGCGGCCCTCTCCCCTTCGCCTTCGACATCCTGGCTACTAGCTTCCTCTACGGCAGCAAGACCTTCATCAACCACACCCAGGGCATCCCCGACTTCTTCAAGCAGTCCTTCCCTGAGGGCTTCACATGGGAGAGAGTCACCACATACGAAGACGGGGGCGTGCTGACCGCTACCCAGGACACCAGCCTCCAGGACGGCTGCCTCATCTACAACGTCAAGATCAGAGGGGTGAACTTCACATCCAACGGCCCTGTGATGCAGAAGAAAACACTCGGCTGGGAGGCCTTCACCGAaACGCTGTACCCCGCTGACGGCGGCCTGGAAGGCAGAAACGACATGGCCCTGAAGCTCGTGGGCGGGAGCCATCTGATCGCAAACATCAAGACCACATATAGATCCAAGAAACCCGCTAAGAACCTCAAGATGCCTGGCGTCTACTATGTGGACTACAGACTGGAAAGAATCAAGGAGGCCAACAACGAGACCTACGTCGAGCAGCACGAGGTGGCAGTGGCCAGATACTGCGACCTCCCTAGCAAACTGGGGCACAAGCTTAATTGAgaattcgatatcaagcttatcggtaatcaacctctggattacaaaatttgtgaaagattgactggtattcttaactatgttgctccttttacgctatgtggatacgctgctttaatgcctttgtatcatgctattgcttcccgtatggctttcattttctcctccttgtataaatcctggttgctgtctctttatgaggagttgtggcccgttgtcaggcaacgtggcgtggtgtgcactgtgtttgctgacgcaacccccactggttggggcattgccaccacctgtcagctcctttccgggactttcgctttccccctccctattgccacggcggaactcatcgccgcctgccttgcccgctgctggacaggggctcggctgttgggcactgacaattccgtggtgttgtcggggaaatcatcgtcctttccttggctgctcgcctgtgttgccacctggattctgcgcgggacgtccttctgctacgtcccttcggccctcaatccagcggaccttccttcccgcggcctgctgccggctctgcggcctcttccgcgtcttcgccttcgccctcagacgagtcggatctccctttgggccgcctccccgcatcgataccgtcgacctcgagacctagaaaaacatggagcaatcacaagtagcaatacagcagctaccaatgctgattgtgcctggctagaagcacaagaggaggaggaggtgggttttccagtcacacctcaggtacctttaagaccaatgacttacaaggcagctgtagatcttagccactttttaaaagaaaaggggggactggaagggctaattcactcccaacgaagacaagatatccttgatctgtggatctaccacacacaaggctacttccctgattggcagaactacacaccagggccagggatcagatatccactgacctttggatggtgctacaagctagtaccagttgagcaagagaaggtagaagaagccaatgaaggagagaacacccgcttgttacaccctgtgagcctgcatgggatggatgacccggagagagaagtattagagtggaggtttgacagccgcctagcatttcatcacatggcccgagagctgcatccggactgtactgggtctctctggttagaccagatctgagcctgggagctctctggctaactagggaacccactgcttaagcctcaataaagcttgccttgagtgcttcaagtagtgtgtgcccgtctgttgtgtgactctggtaactagagatccctcagacccttttagtcagtgtggaaaatctctagca

**Appendix III:**

**mGFP-targeting crispr RNA sequence:**

CTTCAGGGTCAGCTTGGCAG-CGG

**GFP repair oligo template sequence**:

5’-GAACGGCCACAAGTTCAGCGTGTCCGGCGAGGGCGAAGGCGACGCCACATATGGCAAGCTGACCCTGAAGTTCATCTGCACCACCGGCAAGCTGCCCGTG-3’
